# Sociality and nesting strategy shape the bimodal diversity gradient in bees

**DOI:** 10.1101/2025.10.03.680340

**Authors:** Lena R. Heinrich, Aline C. Martins, Silas Bossert, Alice C. Hughes, Michael C. Orr, Katja C. Seltmann, Anshuman Swain, Thais Vasconcelos

## Abstract

Bees are dominant pollinators across native and agricultural plant communities, yet the drivers of their patterns of geographic distribution and functional diversity remain poorly understood. Notably, bees exhibit a bimodal latitudinal diversity gradient, peaking in species diversity in temperate and arid regions rather than the tropics despite their close ecological and evolutionary ties to flowering plants, which show the opposite pattern. Here, we investigated whether two key life history traits thought to shape where bees can live—sociality and nesting strategy—may influence this pattern. We compiled data on sociality and nesting strategy for 4,293 bee species and combined it with a comprehensive phylogeny, curated global occurrence records, and newly-developed models of trait evolution to test 1) whether sociality and nesting biology are evolutionarily correlated, 2) how these life history traits shape bees’ climatic niches and niche breadths, and 3) whether the evolutionary dynamics of these traits may explain the bimodal latitudinal gradient in bee diversity. We find that the evolution of above-ground nesting is closely tied to the evolution of social behavior, but this trait combination has evolved rarely across the evolutionary history of bees. Where this combination does arise, species that are both social and above-ground nesters tend to have narrower climatic niches and are associated with environments that are warmer, wetter, and experience little seasonal temperature variation, making them more prevalent in tropical bee communities. Together, our results indicate that the bimodal diversity gradient in bees is driven by phylogenetic niche conservatism, where most bee lineages retain ancestral traits suited to arid environments and the derived syndrome necessary for thriving in the tropics evolves rarely.

## Introduction

Bees are dominant pollinators of flowering plant communities, and their pollination services and response to global change depend on many functional traits which vary across species, including morphological traits such as tongue length and body size, and behavioral traits such as nesting, foraging, and social behavior (Ostwald et al. 2021; Russell et al. 2024). However, despite bees’ economic and ecological importance across terrestrial ecosystems, we still lack an understanding of where bees with these different functional traits are distributed, what drives the evolution and geographic distributions of these traits, and how they may influence patterns of bee species richness globally (Orr et al. 2021).

Bees are a diverse group of insects including approximately 21,000 described species in seven families, which arose concurrent with the appearance of angiosperms during the Cretaceous (Michener 2007; Almeida et al. 2023). Many bees have specialized interactions with angiosperms, relying on floral resources for food and to provision brood cells for their offspring. As a result, it has been hypothesized that bees have played an important role in angiosperm diversification and vice versa (Michener 2007; Cardinal & Danforth 2013). However, despite their close relationship to angiosperms, bee taxonomists have long hypothesized that bee diversity is low in the tropical latitudes where angiosperm diversity peaks, and highest in xeric and temperate mid-latitudes where angiosperm diversity is relatively low (Michener 1979). This pattern departs from the latitudinal diversity gradient (LDG) where species richness increases from the poles to the tropics, considered one of the most pervasive patterns in ecology (Hillebrand 2004). Recent analyses which mapped the distribution of bee diversity for the first time by synthesizing global bee distribution data sets support this observation (Orr et al. 2021). This bimodal latitudinal diversity gradient in bees, peaking in mid-latitudes, creates a mismatch between bee and angiosperm diversity that is unexpected given their close ecological relationship and raises the question: if not flowering plant diversity, what ecological, environmental, or historical factors drive the spatial distribution of bee diversity?

Since flowering plant diversity and bee diversity appear to be uncoupled, bee diversity patterns may instead be affected by how aspects of bee biology interact with tropical environments, specifically their nesting biology and social behavior. Globally, nearly two-thirds of non-parasitic bee species nest below-ground, constructing brood cells in excavated tunnels or pre-existing cavities and provisioning their larvae with a mixture of pollen and nectar (Cane & Neff 2011). Nesting in the soil likely buffers bees against climatic variation compared to above-ground nests, which are more exposed to fluctuations (Danforth et al. 2019). However, high humidity and precipitation in tropical regions can lead to moisture buildup in brood cells, increasing the risk of fungal growth and spoilage of pollen provisions in below-ground nests (Antoine & Forrest 2021; Potts & Willmer 1997; although some bees are uniquely adapted to seasonally flooded habitats, e.g. see Norden et al. 2003, Roubik & Michener 1980). Nesting above-ground or in cavities, and incorporating waterproofing and antimicrobial materials such as resins, waxes, or floral oils, may mitigate these risks (Buchmann 1987; Chui et al. 2022; Lokvam & Braddock 1999; Vogel 1974). Many tropical bees, including stingless bees (Meliponini), orchid bees (Euglossini), and some oil-collecting bees (e.g. *Centris*), build above-ground nests and use plant-derived materials in their construction (Michener 2007; Martins et al. 2015; Danforth et al. 2019). In *Hoplitis* (Osmiini) bee species, ground-nesters are restricted to xeric and Mediterranean climates and show a limited ability to colonize moister areas (Sedivy et al. 2013), and in the tribe Tapinotaspidini, shifts from the ancestral ground-nesting state to cavity nesting have been concurrent with the colonization of tropical forests (Aguiar et al. 2020), indicating that the environment plays a role in the evolution and distribution of bee nesting strategies.

Social bees may also be more prevalent in the tropics due to more favorable conditions for the emergence of sociality and the potential benefits it offers in these environments, which are characterized by complex, diverse floras and abundant natural enemies. Although the vast majority (∼90%) of non-parasitic bee species are solitary, social taxa are found in several lineages in the families Apidae and Halictidae, encompassing an array of behaviors and sometimes showing a high degree of lability (Cardinal & Danforth 2011; Michener 1969; Peled et al. 2025; Danforth 2002), making bees an ideal group to investigate the evolutionary drivers of sociality. Despite comprising a small fraction of bee species diversity, highly eusocial groups such as stingless bees (Meliponini) and honey bees (Apini) are especially prominent in tropical forests, where they are highly abundant and play an outsized role in structuring pollination networks (Martins et al. 2023; Bueno et al. 2023; Gruchowski-Woitowicz et al. 2024; Ramalho 2004 and citations therein).

Foraging benefits may be one reason for social bees’ prevalence in tropical regions, as social bees’ foraging strategies and colony needs likely align well with tropical floral resource dynamics. Unlike solitary species, most social bees must provision large colonies continuously throughout the year, and consequently require a steady influx of floral resources, foraging opportunistically from a wide range of flowering plants to meet these demands (Devkota et al. 2024; Grüter & Hayes 2022). This strategy is well-suited to the tropics, where floral resources are abundant year-round but patchily distributed across time and space from the perspective of a floral resource specialist. Asynchronous flowering and high flowering plant diversity in the tropics limit the density and continuous availability of any single host, which may pose challenges for specialist foragers that must locate and phenologically track specific host plants (Ogilvie & Forrest 2017; Roubik 1992). For example, among communities of oligolectic bees foraging on creosote bush (*Larrea tridentata*) in the southwestern U.S., specialist bee species richness and abundance are positively correlated with drought frequency, indicating that specialists are favored when flowering phenology is both constrained and highly predictable based on environmental cues like rainfall (Minckley et al. 2000). Together, these attributes may help explain why generalist foragers in the tropics exhibit a greater degree of social behavior.

Furthermore, extended breeding seasons in the tropics may facilitate the evolution of sociality by enabling the production of multiple, overlapping broods—a prerequisite for cooperative brood care and the evolution of complex social behaviors (Kocher et al. 2014; da Silva 2021; Cheng & Ashton 2021).

In addition to foraging advantages, sociality may confer better protection from natural enemies in tropical environments where predation pressures are high, in particular by hyper-abundant, aggressive ants (Jeanne 1979; Roubik 1989, Danforth et al. 2019). During the nest provisioning phase, solitary bees’ larvae are vulnerable to predators that can enter the nest before the brood cell is sealed. In contrast, social bees benefit from a division of labor where some individuals can guard the nest while others forage (Lin & Michener 1972; Evans 1977; Smith et al. 2003; Mikát et al. 2016). In stingless bees, various forms of nest defense have evolved repeatedly in association with the evolution of robber bees, indicating that parasitism risk can drive the evolution of greater social complexity (Grüter et al. 2017). Additionally, sociality in the tropical, facultatively social bee *Megalopta genalis* seems to be driven at least in part by higher rates of nest failure in solitary nests, which are exposed to higher rates of ant predation compared to social nests where the nest is guarded at all times (Smith et al. 2003; Kapheim et al. 2013). In fact, predation pressures are so high in subtropical and tropical environments that trap-nesting studies on bees require the use of sticky materials on supporting stakes to prevent ant predation (Staab et al. 2018).

Together, these ecological perspectives suggest that sociality and nesting strategy may jointly influence where bees can live, and thus may help contextualize global bee diversity patterns. However, the geographic distribution and evolutionary dynamics of these traits remain poorly understood. We lack a clear understanding of where social and above-ground nesting bees are distributed, information necessary to substantiate these hypotheses, and global, phylogenetic analyses that assess whether the evolution of these life history traits are correlated with one another or related to climate. New approaches integrating functional ecology, biogeography, and macroevolution show promise for exploring the drivers of trait-based richness patterns, which can contribute to generating unusual patterns of species richness such as the bimodal diversity gradient seen in bees (Wiens 2023). Here, we use this approach to integrate a newly compiled dataset on sociality and nesting strategy for 4,293 bee species with a comprehensive phylogeny, curated global occurrence records, and recently developed models of correlated trait evolution.

We explore the spatial distribution of social and above-ground nesting bees, and we test: (1) whether sociality and nesting biology are evolutionarily correlated across bees; (2) whether these traits are associated with climate globally; and (3) how these traits influence bees’ climatic niches and niche breadths.

## Methods

### Phylogeny and life history trait scoring

To analyze the evolutionary dynamics of sociality and nesting biology in bees, we used the recently published time-calibrated bee phylogeny from Henríquez-Piskulich et al. (2024) as the basis for all analyses. This phylogeny was inferred using a supermatrix approach, which included both phylogenomic-scale sequence data, as well as single and multi-gene data sets. The topology is highly congruent with comprehensive phylogenomic studies focusing on particular groups of bees (e.g., Bossert et al. 2022; Sless et al. 2022; Freitas et al. 2020), and the inclusion of 4,586 species (∼23% of all described bee species and 82% of genera) renders this tree a sound basis for testing the evolution of traits from a phylogenetic perspective.

All species represented as tips in this tree were scored with respect to nesting biology (below-ground or above-ground) and sociality (simplified into solitary or social for trait evolution analyses; see trait scoring details below) through a literature survey updated through August 2024. First, we scored broad taxonomic groups with consistent nesting or social behavior using Michener (2007)–for example, the families Andrenidae, Megachilidae, Melittidae, Stenotritidae, and Colletidae (except *Amphylaeus morosus*, see Spessa et al. 2000) do not include any social species, as far as is currently known. For species in groups with variable nesting or social behavior, we performed individual Google Scholar searches for each taxon using first the genus name and then the full species name with the keywords “nest”, “nesting biology”, “social”, and “solitary”. For example, we queried Google Scholar using the search string “Xylocopa” AND (“sociality” OR “social” OR “solitary”). If results indicated consistent social behavior across the genus, we scored all species accordingly; if behavior was variable, we repeated the search using the full specific epithet.

Bee nesting behavior is diverse, but can be broadly divided by nest location into above- and below-ground. The majority of bees (64% – 83%) nest beneath the soil surface, either by excavating their own burrows or by using those left by rodents (Antoine & Forrest 2020), as is common in the genus *Bombus*. Above-ground nesting includes both cavity and aerial nesters. Cavity nesters exploit a range of substrates, including dead wood, hollow stems, crevices in stones or trees, empty shells, or abandoned nests of other insects (Michener 2007; Danforth et al. 2019), whereas aerial nesting is restricted to highly eusocial bees such as honey bees (*Apis*) and some stingless bees (Meliponini), which construct exposed nests *de novo* out of waxes and resins (Michener 1974; Grüter 2020).

Similarly, social behavior in bees is continuous, ranging from completely solitary to highly eusocial with various intermediate stages, including communal, parasocial, primitively eusocial, and highly eusocial behavior (Michener 1969; Peled et al., 2025). Most bees are solitary (∼90%), although some form aggregations, nesting separately but in close proximity, or nest communally, sharing a nest entrance but provisioning offspring independently (Michener 2007). Parasocial bees share a nest, and may cooperate in brood care with some division of labor, but lack distinct castes and typically involve females of the same generation (in contrast to the matrifilial colonies of eusocial bees). Primitively eusocial bees, including many bumble bee (*Bombus*) species, are matrifilial with cooperative brood care and division of labor, but have poorly differentiated castes and a solitary nest-founding phase. Finally, highly eusocial bees like *Apis* and Meliponini have all of the traits of primitively eusocial species but maintain social colonies year-round with clearly defined castes (Michener 1969).

Although communal or aggregate nesting has been observed in many species, it is poorly established how commonly these aggregations form and whether they are social in nature or a consequence of common nesting habitat selection among conspecifics (Michener 1969). To include only species with definitive social behavior, species were only scored as social if they displayed consistent parasocial behavior or above (i.e., some degree of cooperative brood care or overlapping generations). This binary coding is a necessary simplification to ensure that only species with well-established social behavior are treated as social, since the degree of sociality remains poorly documented for many species and existing categories are used inconsistently across the literature, making it difficult to ensure that more nuanced classifications represent cohesive and comparable units. Likewise, nesting categories were simplified to “ground” or “above-ground” since above-ground nesters can often utilize a diverse range of substrates which are likely to be similar in their exposure to the abiotic environment relative to nesting in the ground. Collapsing the categories into two binary traits also reduces the number of parameters to be estimated and increases the statistical power of trait evolution analyses.

Polymorphic species (i.e., those that are facultatively social or which will flexibly nest either below or above-ground) or species which lacked reliable natural history information were excluded from analyses (245 species in total). Additionally, all brood parasites were scored as solitary and brood parasite nesting behavior was scored based on that of their hosts (i.e., a brood parasite which parasitizes a ground-nesting bee would also be scored as ground-nesting, rather than as a separate brood parasite category). Because brood parasites closely associate with their hosts and do not construct their own nests, we reason that their relationship with climate will be driven more by their host species than by the state of being parasitic. Similarly, social parasites such as some *Lasioglossum* and *Bombus* species were scored as social and their nesting behavior scored on the basis of their host.

### Climatic niche characterization

To characterize species’ climatic niches, bee occurrence points were obtained from Dorey et al. (2023), which compiles data from large aggregators such as GBIF, SCAN, iDigBio, USGS, and ALA. We then cleaned the data using the *BeeBDC* package in R by following the full occurrence point cleaning workflow (Dorey et al. 2023). The resulting dataset included 6,890,148 occurrence points for 11,607 bee species. After subsetting this dataset to include only species also present in the Henríquez-Piskulich et al. (2024) phylogeny, 6,128,948 occurrence points remained. This included 3,952 of the original 4,586 species present in the phylogeny, indicating 634 species lacked occurrence data that met the criteria used in the *BeeBDC* filtering process (Dorey et al. 2023). These species were excluded from trait-environment correlations and niche breadth analyses, in addition to nineteen species known to have been introduced outside of their native ranges (Goulson 2003; Table S1). To reduce biases from oversampling in some regions, we thinned the remaining occurrence records to a maximum of three points per species within each one-degree grid cell, and then overlaid them with 19 bioclimatic layers from the CHELSA (“Climatologies at high resolution for the earth’s land surface areas”) dataset (Karger et al. 2017). Although global bioclimatic datasets like CHELSA have some limitations such as interpolation errors in data-poor or topographically complex regions, CHELSA represents one of the highest-resolution and most robust datasets currently available. Climate values were extracted for each occurrence point using the R packages *sp* and *raster* and averaged for all occurrence points per bee species to estimate their climatic niche (Pebesma & Bivand 2005; Hijmans 2022; pipeline adapted from Vasconcelos et al. 2023). Finally, we performed Kruskal–Wallis tests with Bonferroni-corrected Dunn’s post hoc comparisons to characterize the distribution of mean climatic variables for each trait combination and document geographic variation in bee community composition. However, because these analyses do not account for phylogenetic relatedness (Felsenstein 1985), their results should be interpreted only as descriptive patterns.

### Mapping the spatial distribution of sociality and nesting strategies

If sociality and nesting strategy shape the spatial distribution of bee diversity, we predict that social, above-ground nesters should make up a higher proportion of the bee fauna in tropical regions, where overall bee species richness is low. To visualize this pattern, we mapped proportions of social bees and above-ground nesting bees separately across a latitudinal gradient in the Americas, where occurrence data for bees is the most available (Orr et al. 2020), to evaluate geographic trends in their distributions. Using occurrence data from Dorey et al. (2023) for species we had scored for sociality and nesting biology, we calculated total bee species richness, as well as the proportion of social and above-ground nesting species, within each polygon defined at level 3 of the World Geographical Scheme for Recording Plant Distributions (TDWG 2001). These polygons correspond roughly to political boundaries, with some larger regions further divided into states. To explicitly visualize how trait prevalence varies with latitude, we then generated scatterplots showing the proportion of species that are social or above-ground nesting as a function of absolute latitude, where each point represents a polygon and is size-scaled by total richness of bee species present in our dataset. To address potential spatial autocorrelation and account for nonlinear relationships between trait proportions and latitude, we fit generalized least squares models with quadratic terms for absolute latitude and a spherical spatial correlation structure.

### Macroevolutionary dynamics of sociality and nesting strategies

Because sociality and nesting strategy may jointly influence bees’ ecological tolerances and evolutionary trajectories, we treat these traits as combined syndromes for analyses. Specifically, we examine four combined states: solitary + ground nesters, solitary + above-ground nesters, social + ground nesters, and social + above-ground nesters (hereafter, *sociality-nesting strategies*). Additionally, we hypothesized that nesting biology and sociality would be evolutionarily correlated, such that social bees are more likely to nest above-ground and solitary bees below-ground, and that these combinations would be associated with two extremes in the climatic gradient: social + above ground nesters with warmer, wetter, less seasonal environments, and solitary + ground nesters with cooler, drier, and more seasonal environments.

To evaluate whether sociality and nesting strategy are correlated, we used the R package *corHMM*, which implements models of discrete character evolution, to estimate transition rates between sociality-nesting strategy states. Using the *fitCorrelationTest* function, we compared four models: (1) an independent model assuming no correlation between traits, (2) a correlated model allowing joint evolution of traits, and (3–4) their respective Hidden Markov Model variants, which account for rate heterogeneity across the tree (Boyko and Beaulieu 2023). In these models, dual transitions are disallowed, both because it is highly unlikely that both traits (i.e., sociality and nesting) would change simultaneously and because assuming stepwise transitions is necessary to test for dependence in discrete trait correlation (Pagel 1994). The Hidden Markov model approach is particularly powerful for large, ancient clades like bees because it does not assume a single, constant rate of trait evolution across the phylogeny. Instead, it introduces “hidden states” that reflect differences in evolutionary rate, allowing for different parts of the tree to have “fast” or “slow” rates of evolution. This better reflects the biological reality that some lineages are more evolutionarily labile than others, improving the accuracy of ancestral state reconstructions (Beaulieu et al. 2013; King & Lee 2015). Support for either of the correlated models (with or without hidden states) would suggest that sociality and nesting strategy do not evolve independently. Finally, we used the recently-developed *corHMMdredge* approach (Boyko 2025) to identify the best-fitting model for transitions and transition rates between sociality–nesting strategies. This model also provided the transition rates used in our ancestral state reconstructions and visualizations of trait evolution across the bee phylogeny.

For ancestral state reconstructions, we set the root state to solitary and ground-nesting based on previous evidence from both bees and their putative sister group (Sann et al. 2018; Debevec et al. 2012; Branstetter et al. 2017; Michener 1974, 2007). Sann et al. (2018) identified Ammoplanidae as the closest extant relatives of bees, which, based on morphology, consists primarily of solitary ground-nesting wasps (Krombein 1979). Together with the observation that the early-branching bee family Melittidae shares these traits (Michener 2007; Danforth et al. 2013), these findings strongly support that the most recent common ancestor of all bees was also solitary and ground-nesting. Finally, we combined the different rate classes under the observed state for ancestral state reconstructions and used the best-fit *corHMM* model to run 100 stochastic maps using the function *makeSimmap* to summarize the number and ages of evolutionary transitions.

### Evolutionary associations between sociality, nesting strategy and climatic niche

To model the evolution of climatic niche in relation to sociality and nesting strategy, we used the function *hOUwie* in the R package *OUwie* (Beaulieu et al. 2012; Boyko et al. 2023), which jointly models discrete and continuous trait evolution under an Ornstein-Uhlenbeck (OU) process, in this case sociality-nesting strategy and climatic niche respectively. By integrating discrete and continuous trait evolution and accounting for heterogeneity in evolutionary rates, *hOUwie* can allow for a more nuanced understanding of how climatic niche evolution may be shaped by, and co-evolve with, key ecological traits (Boyko et al. 2023).

We used the best-fitting model from the *corHMMdredge* analyses as the base model for mapping discrete character states (i.e., sociality–nesting strategies corresponding to distinct evolutionary regimes) across the phylogeny, and for estimating the climatic niche optimum (θ) associated with each regime. In our case, θ represents how bees with different sociality-nesting strategies differ in their climatic optima over time. We ran six versions of the continuous trait model to test different hypotheses about how climatic niche might evolve across sociality-nesting strategies: (1) a Brownian motion model assuming a single rate of evolution across the tree, (2) an OU model with a single climatic optimum for all species, (3–4) OU models where optima differed depending on sociality (3) or nesting strategy (4), (5) an OU model with a distinct optimum for each of the four discrete trait combinations, and (6) a character-independent OU model in which climatic optima can differ between hidden rate classes but are shared across observed states, so trait identity does not influence the optimum beyond the rate class. Each model was run with 100 simulations and 10 starting points. We ran these models separately for each of four focal climatic variables: mean annual temperature (BIO1), temperature seasonality (BIO4), annual precipitation (BIO12), and precipitation seasonality (BIO15), chosen to summarize both temperature and precipitation variables and their seasonal variation.

### Comparing climatic niche breadths across sociality-nesting strategies

Finally, to evaluate how sociality and nesting strategy shape bees’ climatic niche breadths, we quantified both univariate and multivariate climatic niche breadths for each sociality-nesting strategy.

For univariate niche breadths, we calculated temperature and precipitation ranges for each species using species-level climatic means from global occurrence records. Temperate breadth was calculated as the difference between the maximum temperature of the warmest month (BIO5) and the minimum temperature of the coldest month (BIO6), and precipitation breadth as the difference between precipitation of the wettest month (BIO13) and precipitation of the driest month (BIO14). Differences among sociality–nesting strategies were then tested using a permutational multivariate analysis of variance (PERMANOVA) with the *adonis2()* function from the package *vegan* based on Euclidean distances and 999 permutations (Oksanen et al. 2025), followed by pairwise comparisons.

For multivariate niche breadths, we performed a principal component analysis (PCA) on the 19 bioclimatic variables from CHELSA (Karger et al. 2017), after excluding highly correlated variables (Pearson |r| > 0.9) and standardizing all variables (mean = 0, SD = 1). The first three principal components were retained to represent climatic niche space, and we used 3D alpha hulls calculated using the *alphashape3d* package to estimate the volume of this space occupied by each sociality–nesting strategy (Lafarge et al. 2023). Pairwise differences in multivariate climatic niche space were tested with PERMANOVA using the same parameters as above. All code and data needed to reproduce these analyses are available at https://github.com/tncvasconcelos/bee_sociality_dist.

## Results

### Trait dataset and phylogeny

Our nesting and sociality trait dataset encompassed 4,293 species, and recovered most species as solitary (75%) and ground-nesting (68%), with 24% being social and 31% above-ground nesting species (Table S2). Solitary ground-nesting species comprised about half of the dataset (51%), while social above-ground nesters were the least numerous trait combination (7.55%). Three families did not vary in sociality or nesting behavior (Andrenidae, Melittidae, and Stenotritidae), while others were more variable (e.g. Apidae, which encompassed all four possible state combinations, Colletidae and Halictidae, which encompassed three states, and Megachilidae, which encompassed two states) (Figure S1).

### Where do social and above-ground nesting bees live?

Mapping of bees’ sociality and nesting strategies across the Americas revealed that the proportion of social bees was highest in boreal Greenland and northern Canada, where the social genus *Bombus* is the most common or only bee taxon present, and also elevated around tropical northwest and central South America. In contrast, solitary bees comprised the majority of bee species diversity in the western United States, Argentina, and Chile (Figure 2A). Above-ground nesting bee species were prevalent across Central America and in northwest and central South America, in particular around the Amazon basin. Ground-nesting bees predominated outside of these regions (Figure 2C).

**Figure 1.**
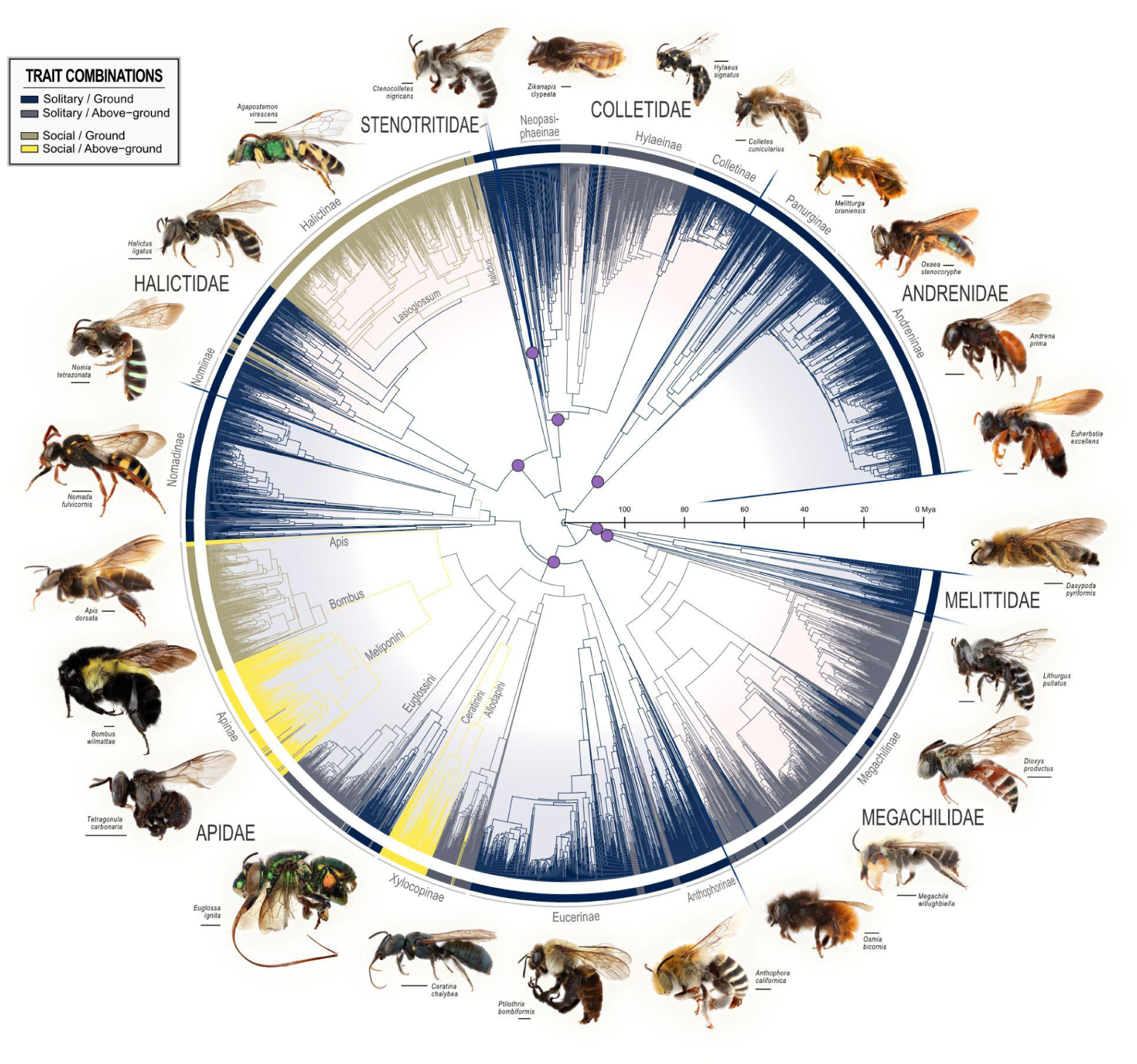
Maximum likelihood phylogeny of bees from Henríquez-Piskulich et al. (2023), pruned to include only species with complete trait data for both sociality and nesting biology (*n* = 4,293). Trait combinations are mapped along the tips and branches are colored based on the most likely ancestral state for that branch when marginal likelihood is ≥ 0.5 (otherwise colored light gray). Ancestral state reconstructions were inferred using a hidden rates model in corHMM with two rate classes and a fixed root state (solitary/ground). Internal purple dots denote the most recent common ancestor of each family.

**Figure 2.**
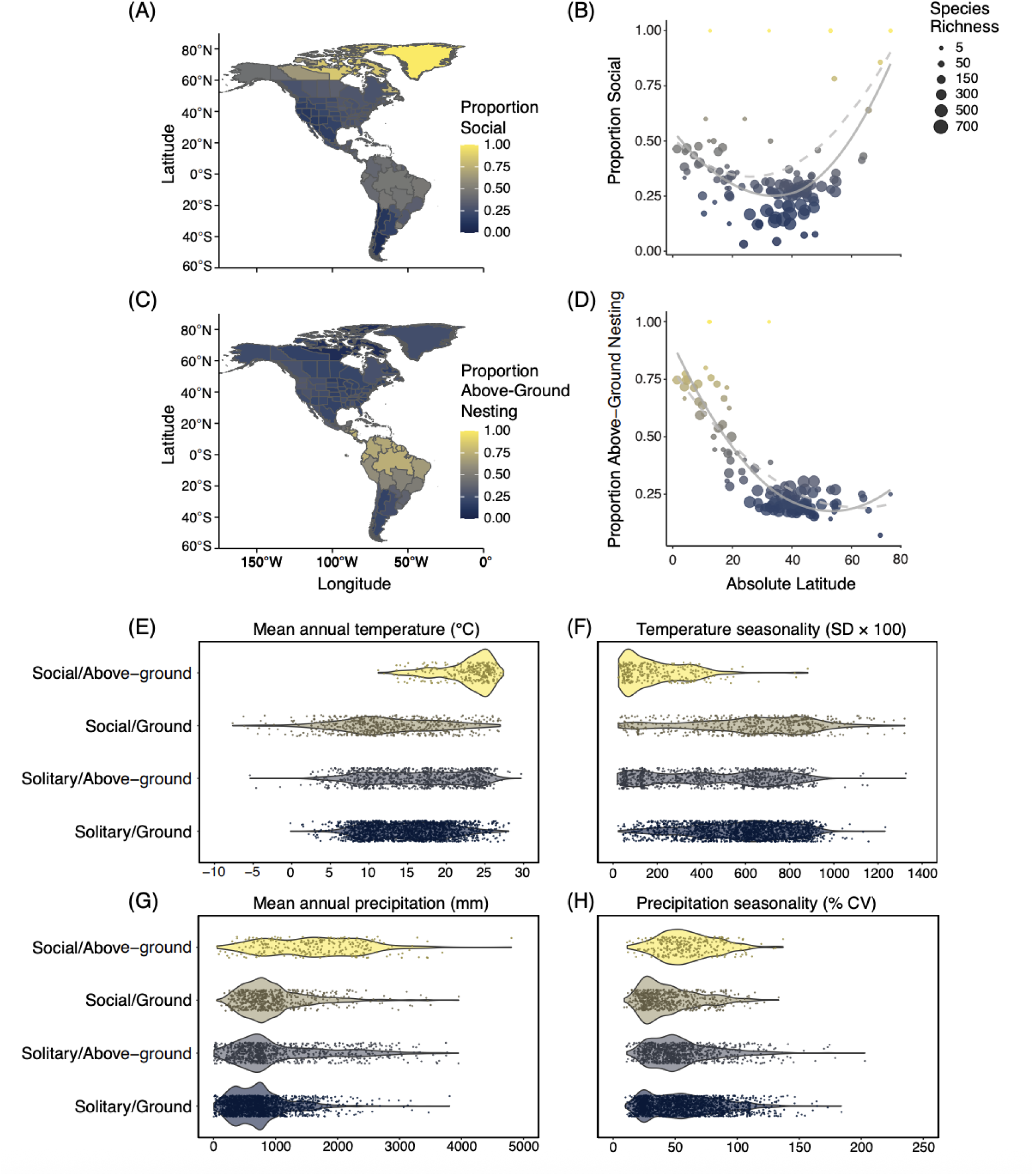
Proportion social **(A)** and proportion above-ground nesting **(C)** bee species relative to total bee species richness across the Americas, and the same data visualized as scatterplots of proportion social **(B)** and proportion above-ground nesting **(D)** species with respect to absolute latitude. Scatterplot points represent regions, with their position showing the trait proportion and their size reflecting total bee species richness in our dataset. Trendlines were fit using generalized least squares models with quadratic terms for absolute latitude, and dashed trendlines are corrected for spatial autocorrelation. Total species richness was tallied by counting unique bee species present in each area defined by TDWG’s level 3 botanical countries using bee occurrence points from Dorey et al. (2023) for species we had scored for sociality and nesting biology. Proportion social and proportion above-ground nesting species were calculated by tallying the number of social and above-ground nesting species relative to this total richness. **(E-H)** Distribution of bees with different sociality-nesting strategies across focal climate variables using global occurrence data (*p* < 0.0001 for all pairwise comparisons using Kruskal-Wallis test).

Scatterplots revealed non-linear latitudinal structuring which was robust to spatial autocorrelation for both sociality and nesting strategy, but in different ways (Figure 2B, D). The proportion of social species exhibited a U-shaped relationship with absolute latitude, with higher proportions of social species both near the equator and at high latitudes, and the lowest proportions observed at mid-latitudes (∼30–50°) (Figure 2B). In contrast, the proportion of above-ground nesting species declined sharply with increasing latitude, with high proportions at low latitudes followed by a sharp drop beyond ∼20–30° after which proportions remained consistently low, indicating a strong association between above-ground nesting and low latitudes (Figure 2D). These geographic patterns were consistent with nonparametric comparisons across trait combinations, where Kruskal–Wallis tests with post hoc contrasts showed that species which were both social and above-ground nesting tended to occur in warmer, wetter regions characterized by reduced temperature seasonality and increased precipitation seasonality relative to other trait combinations (Figure 2E-H; see Table S3 and Figure S2 for analysis of all climate variables).

### Are sociality and nesting strategy evolutionarily correlated?

The *corHMM* analyses indicated that sociality and nesting strategy have evolved in a correlated manner in bees. The best-supported model was the hidden Markov correlated model (Table 1), indicating that transitions in these two traits are not independent from each other and are best described by a model allowing for rate heterogeneity across the tree. Specifically, the best fit model from the *corHMMdredge* framework includes a “fast” rate class (R1), where transitions between trait states occur more frequently, reflecting greater lability, and a “slow” rate class (R2), where transitions are less frequent, reflecting stronger conservation of the ancestral state. Under this model, the root state was inferred as being rate class 1 (R1) solitary + ground nesting, and the most probable transitions overall were between R1 and R2 within the solitary + ground nesting state, or reverting from a solitary + above-ground nesting state to the ancestral state of solitary + ground nesting in the fast rate class, R1 (both 0.041 events per million years). This was far more likely than reverting from above-ground to ground nesting as a social bee (0.008 events per million years), suggesting greater evolutionary lability in nesting strategies among solitary bees. Conversely, in the slow rate class (R2), no reversals from above-ground to ground nesting were detected for either solitary or social bees, implying unidirectional shifts to above-ground nesting in lineages under this rate class (Figure 3A).

**Figure 3.**
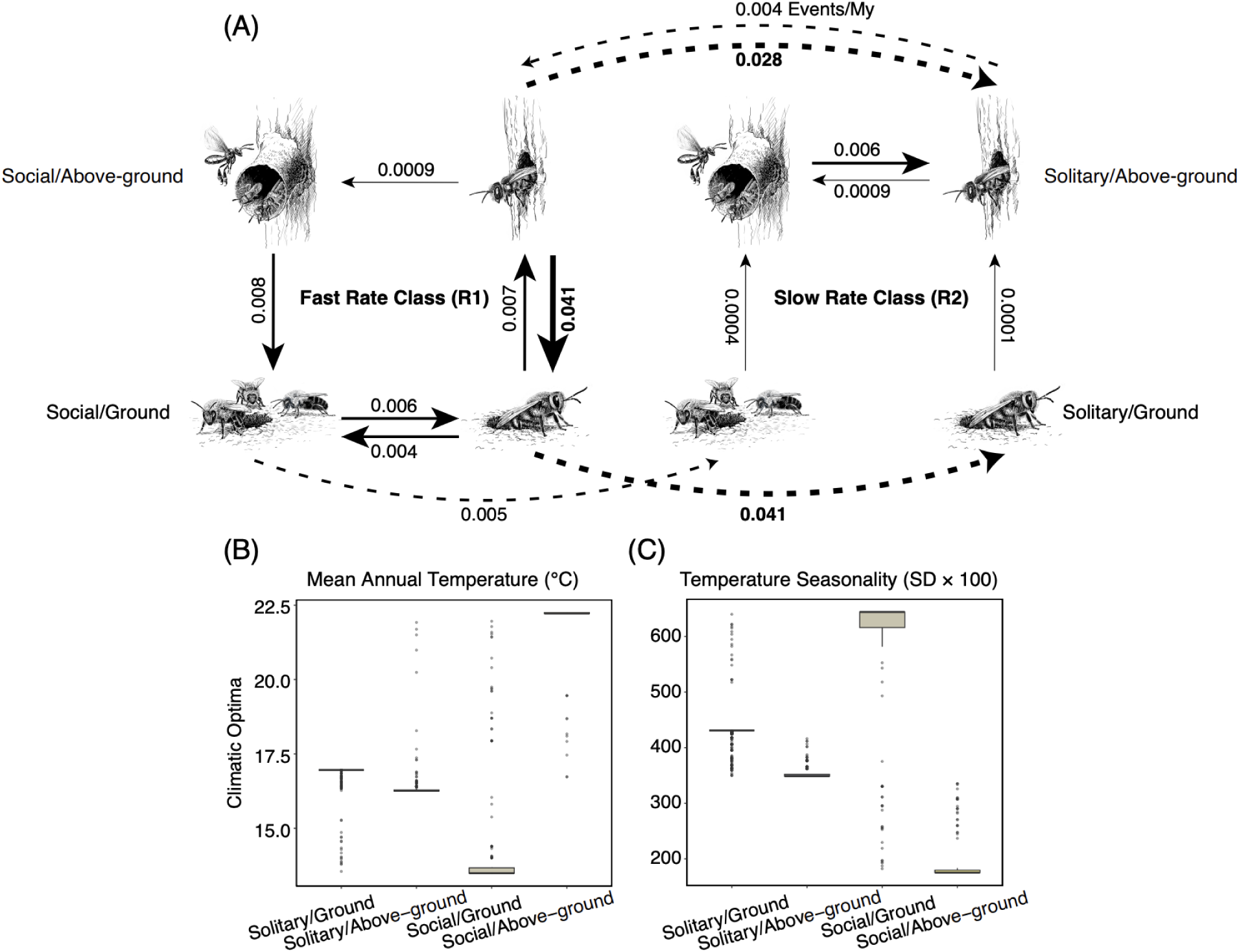
(A) Evolutionary transition rates between sociality-nesting strategies in bees. Transition rates (events per million years) were estimated using *corHMMdredge*, based on the best-fitting correlated Hidden Markov model which includes two rate classes: a fast rate class (R1), where trait evolution occurs more quickly, and a slow rate class (R2), where trait evolution occurs more slowly. Arrows represent transition rates between states, with solid lines indicating transitions between character states and dashed lines indicating transitions between rate classes. Arrow widths are scaled to estimated rates. Illustrations by John Megahan (University of Michigan Museum of Zoology). **(B-C)** Distribution of model-averaged expected means (i.e., estimates of the evolutionary climatic optima, or θ) inferred from hOUwie models for each combination of sociality and nesting biology across **(B)** mean annual temperature (BIO1) and **(C)** temperature seasonality (BIO4).

**Table 1.** Comparison of corHMM models evaluating the evolution of sociality and nesting strategy in bees using a fixed root state of solitary and ground-nesting. Models differ in their assumed patterns of trait change (independent vs. correlated evolution) and the number of hidden rate classes (1 or 2).

| Model | np | lnLik | AIC | ΔAIC | AIC Weight |
| --- | --- | --- | --- | --- | --- |
| **Hidden Markov Correlated** | **18** | **-544.69** | **1125.39** | **0.00** | **1.00** |
| Hidden Markov Independent | 10 | -579.10 | 1178.20 | 52.81 | $3.41 \times 10^{-12}$ |
| Correlated | 8 | -604.62 | 1225.24 | 99.85 | $2.08 \times 10^{-22}$ |
| Independent | 4 | -616.53 | 1241.06 | 115.67 | $7.61 \times 10^{-26}$ |

Transitions to the social + above-ground state were consistently rare across both rate classes, with estimated rates of just 0.0009 events per million years from solitary + above-ground in both the fast and slow rate classes (R1 and R2), and less than half that likelihood—0.0004 events per million years—from social + ground in the slow rate class (R2). This asymmetry suggests that sociality is more likely to evolve after a lineage has already adopted above-ground nesting, rather than from a ground-nesting ancestor. Although transitions from a solitary to a social state occurred more frequently among ground-nesting taxa (0.004 events per million years), subsequent shifts to above-ground nesting were either very unlikely (R2) or not observed (R1) once sociality had already evolved (Figure 3A). The most probable evolutionary pathway to a social + above-ground state, then, appears to be a stepwise transition: from the ancestral solitary + ground-nesting condition, first to solitary + above-ground, and last to social + above-ground. This requirement for a stepwise transition, coupled with the rarity of such transitions, has made the evolution of the social + above-ground nesting combination exceedingly rare across the evolutionary history of bees, with an average of 15 transitions to this state, all within Apidae (see Figures S3 & S4, Table S4 for a summary of all transitions).

### Is bee climatic niche evolution associated with sociality and nesting biology?

Climatic niche evolution in bees was associated with sociality and nesting strategy for temperature-related variables, but not for precipitation or precipitation seasonality. For both mean annual temperature (BIO1) and temperature seasonality (BIO4), the best-supported models assumed different climatic optima for each of the four sociality-nesting strategies, with strong support (AICc weight > 0.97). In contrast, annual precipitation (BIO12) was best explained by a model that allowed optima to vary freely across observed states without tying them to specific traits (AICc weight = 0.995), and precipitation seasonality (BIO15) was best explained by a model where all sociality-nesting strategies had the same optima (AICc weight = 0.9987) (Table S5).

Social + above-ground nesting bees occupied niches with the highest mean annual temperature (∼22.1 °C), while social + ground nesting bees occupied the coolest (∼13.8 °C). Solitary strategies fell in between (∼16.3–16.9 °C). Similarly, social + above-ground nesting bees were associated with environments with the least temperature seasonality (∼186 SD × 100), and social + ground nesting bees with the most seasonal (∼618 SD × 100), with solitary strategies again intermediate (Figure 3B-C; Table S6). Together, these results indicate that evolutionary shifts in thermal environments appear to be structured by sociality and nesting biology in bees, with social + above-ground nesting bees occupying warmer environments with less seasonal variation in temperature.

### How do sociality and nesting strategy influence bees’ climatic niche breadths?

We found strong differences in both univariate and multivariate climatic niche breadth among the four bee sociality-nesting strategies. In univariate comparisons, species classified as social + above-ground nesters exhibited the narrowest temperature and precipitation niche breadths, while the other three groups (solitary + ground-nesters, solitary + above-ground nesters, and social + ground-nesters) occupied broader ranges of both temperature and precipitation (Figure 4B-C, Table S7).

**Figure 4.**
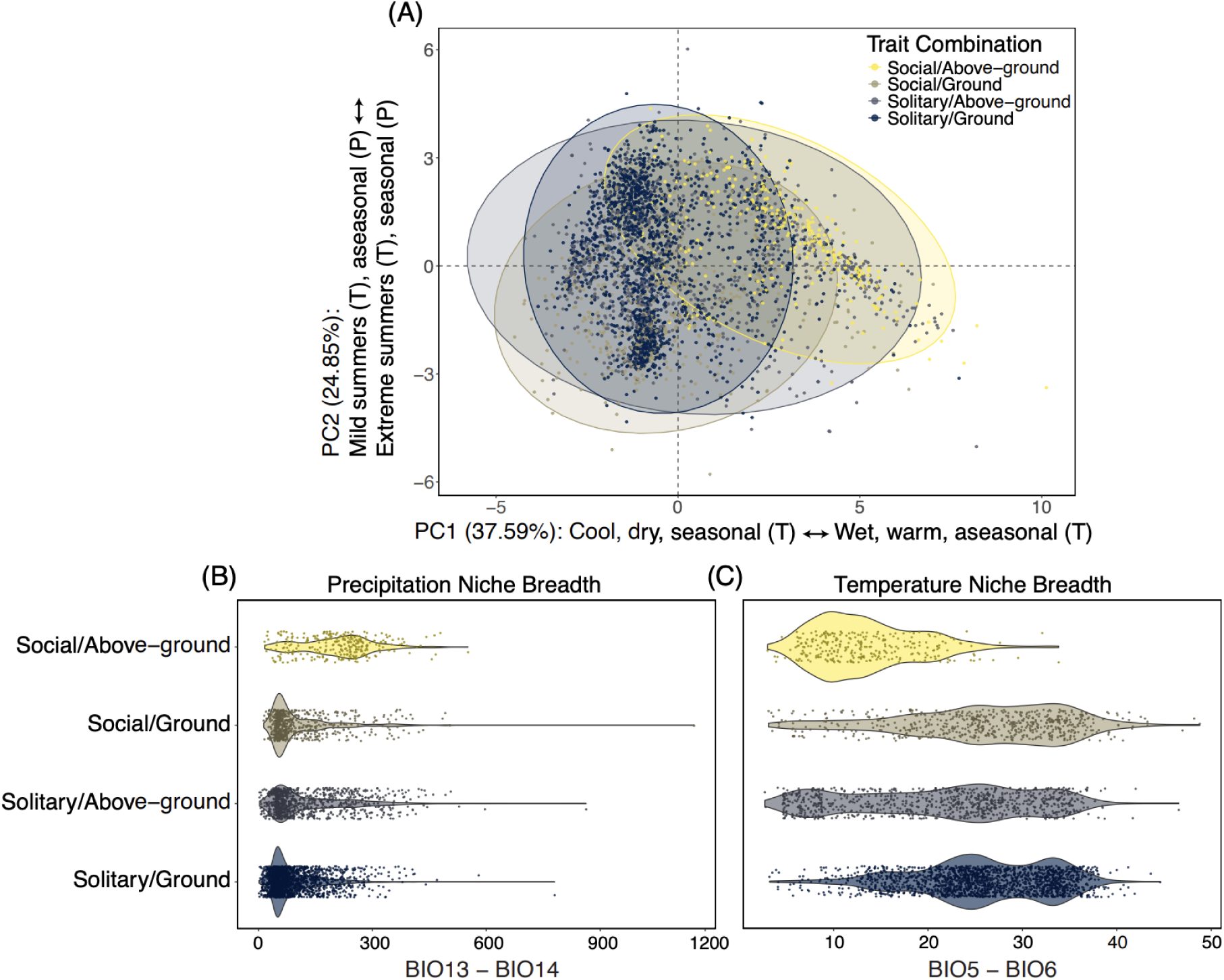
**(A)** Principal component analysis (PCA) of multivariate climatic niche space across bee species using two principal components comparing sociality-nesting strategies. Each point represents a species’ position in the climatic niche space based on 12 uncorrelated (|r| < 0.9) and standardized (mean = 0, SD = 1) bioclimatic variables, reduced using PCA. Points represent individual bee species, and shaded polygons are 95% normal-distribution ellipses for each trait combination. PC1 and PC2 explain 37.59% and 24.85% of the total climatic variation across species respectively, for a total of 62.44% together. PC axis titles were characterized based on the PCA loadings and contributions: higher scores for PC1 indicate higher mean annual temperatures, more rainfall overall, especially in the wettest and warmest quarters, and lower temperature seasonality (“aseasonal (T)”, more thermally stable climates), whereas higher scores for PC2 indicate higher precipitation seasonality (“seasonal (P)”, distinct wet and dry periods), higher maximum temperatures in the warmest month, and greater daily temperature variation (Figure S5). Social + above-ground nesting bees were primarily distributed on the right side of the PCA, corresponding to higher values of PC1 or warmer, wetter environments with less seasonal variation in temperature **(B-C)** Univariate climatic niche breadths (precipitation, **B**, and temperature, **C**) across bee species with different combinations of sociality and nesting strategy. Niche breadths were calculated as the difference between temperature and precipitation extremes across occurrence points.

Principal component analysis revealed that the first three axes accounted for 78.5% of total climatic variation. These components were used to construct a multivariate climatic niche space in which each species was positioned. Volumetric analyses of this space showed that social + above-ground nesters occupied the smallest multivariate niche volume (68.1 units), while the remaining groups occupied significantly larger and more similar volumes: 254 (solitary + above-ground), 250 (social + ground-nesting), and 242.4 (solitary + ground-nesting) units (Table S8, visualized using two principal components in Figure 4A). A global PERMANOVA indicated that trait groups differ significantly in their positions in multivariate climate space (F = 160.7, R² = 0.115, p = 0.001), with sociality-nesting strategy explaining 11.5% of the variance in climatic niche position across species. Pairwise PERMANOVAs revealed that all groups occupy significantly different positions in the climatic niche space (*p* = 0.001 for all), though the magnitude of separation varied. The greatest niche divergence occurred between social + above-ground and social + ground-nesting species (R² = 0.255), followed by the contrast between social + above-ground and solitary + ground-nesting species (R² = 0.165). Comparisons among solitary groups, or between solitary and social ground-nesting species, yielded lower R² values (∼0.03), indicating relatively higher niche overlap among these strategies (Figure 4A, Table S9).

## Discussion

Together, our analyses indicate a complex ecological and evolutionary interplay among sociality, nesting strategy, and geographic distribution in bees. We find that the solitary, ground-nesting state dominates in arid and seasonal regions; the solitary, above-ground nesting state occurs in similar, slightly broader climatic contexts; the social, ground-nesting state is concentrated in cool, seasonal environments such as those occupied by bumble bees (*Bombus*); and, finally, the social, above-ground nesting state occupies the narrowest climatic niche centered over environments that are warm, wet, and with little seasonal variation in temperature. Trait evolution analyses revealed that sociality and nesting strategies are evolutionarily correlated, indicating that these life history traits may arise in response to shared ecological pressures, particularly in relation to temperature variables. Finally, our results suggest that the combination of sociality and above-ground nesting has evolved infrequently across the bee phylogeny. Low transition rates, combined with the requirement for a two-step transition from the ancestral state of solitary and ground nesting, may help explain why social above-ground nesting bees make up only a small fraction of modern global bee diversity despite being prominent in tropical environments. Furthermore, these evolutionary constraints offer a compelling explanation for the unexpectedly low diversity of tropical bee faunas, since the traits which appear to be most conducive to persistence in those environments evolve rarely.

### Social, above-ground nesting strategy dominates among tropical bees

Although both sociality and above-ground nesting have been individually hypothesized to be advantageous under tropical conditions, our results show that the combination of these traits may act synergistically, structuring bee distributions more than either trait independently and partitioning bees into distinct ecological regimes. Social, above-ground nesting bees, in particular, are disproportionately represented in tropical communities. This category is skewed towards two closely related, speciose, highly eusocial clades (i.e. Apini and Meliponini) that often form large, perennial colonies, features that could help explain their success in tropical environments by enhancing defense and foraging capacity (Michener 1974, Roubik 1989, Roubik 2023). Bumble bees, which remain ground-nesting despite being social, are an informative exception that illustrate the constraints faced by other social lineages in cold, seasonal environments. Unlike many highly eusocial tropical species, bumble bee colonies are usually small and annual rather than perennial, with queens entering diapause during resource-poor winter months (Michener 1974), although some tropical bumble bee species can maintain perennial colonies (Cameron & Jost 1998, Oliveira et al. 2022). Their flexible generation times allow them to match resource demands with floral availability, a strategy that may support persistence in temperate and boreal regions where floral resource availability fluctuates (Ogilvie & Forrest 2017). In addition, bumble bees possess unique thermoregulatory abilities that enable them to forage and maintain colony activity under cool conditions (Heinrich 1972, 1975). These adaptations may explain why bumble bees are the primary radiation of social bees in cold, highly seasonal regions where most other social lineages have failed to establish as part of their native range.

While the combination of sociality and above-ground nesting appears ecologically significant, our results suggest that nesting strategy exerts a stronger influence on climatic niche differentiation than sociality. Climatic niche divergence was most pronounced between above- and ground-nesting species, regardless of sociality, indicating that nesting strategy is the primary axis structuring climatic niches in bees. Our spatial mapping of life history strategies across the Americas also supports this interpretation: nesting strategy is more strongly associated with latitude than sociality is, as illustrated by the prevalence of social, ground-nesting bumble bees (*Bombus*) at high latitudes (Williams 1998; Hines 2008).

The transition from ground-nesting to above-ground nesting likely represents a major ecological shift in the evolution of bees, decoupling species from soil microclimates and the community of soil-associated predators, pathogens, and competitors that are especially active in warm, stable environments (Danforth et al. 2019; Wcislo 1987). In this sense, shifts in nesting strategy may enable subsequent shifts in social organization by reducing exposure to these selective pressures, explaining why the evolution of above-ground nesting often precedes the evolution of sociality in lineages with this trait combination. The association between nesting substrate and social organization may also reflect a functional link between above-ground nesting and the evolution of large, perennial social colonies (da Silva 2021). Above-ground nesters can exploit a wide range of pre-existing cavities or build aerial nests tailored to colony size, whereas ground-nesting species are constrained by the energetic costs of excavation and the limited sizes of available burrows. This connection is again exemplified by bumble bees, whose ground-nesting habit coincides with smaller, annual colonies (Michener 1974), suggesting that nesting substrate may impose an upper limit on colony size and the level of social organization that can be attained. Together, these results suggest that nesting strategy may mediate the link between climate and social evolution, with shifts to above-ground nesting opening new climatic space that facilitates the origin and persistence of highly social lineages in the tropics by enabling the development of larger, more defensible colonies with greater foraging capacity.

### The evolution of sociality and nesting strategy in bees is linked to temperature

Although aspects of bees’ biology suggest that both temperature and precipitation may shape the evolution of sociality and nesting strategy, we find that evolutionary divergence in climatic niches across trait combinations is primarily associated with temperature-related variables rather than precipitation. Phylogenetic analyses using *hOUwie* indicate that only temperature-related variables (mean annual temperature and temperature seasonality) show evidence of evolutionary divergence aligned with sociality and nesting strategy. Specifically, species with different sociality-nesting strategies have different thermal optimas, with social + above-ground nesting bees evolving in habitats where temperatures are warmer and less variable across the year, and social + ground nesting bees in cooler environments that experience greater seasonal temperature fluctuations. This pattern aligns with the idea that nesting below-ground buffers bees from climatic extremes, whereas above-ground nests are more exposed to fluctuations and therefore associated with stable environments (Danforth et al. 2019). Moreover, because latitude is more closely tied to temperature than to precipitation, the association of nesting strategy with latitude also reflects the importance of thermal regimes in structuring where above-versus below-ground nesting species can thrive. Overall, temperature appears to have been more important than precipitation in shaping trait-associated climatic niches over evolutionary time, while the observed association between sociality, above-ground nesting, and wetter environments may be mediated by other processes.

One possibility is that a clearer evolutionary signal with precipitation could emerge with more refined trait categories—for example, distinguishing social lineages by colony size, annual versus perennial life cycles, or by using eusociality as the cutoff for sociality—since these distinctions may interact more directly with rainfall stochasticity. However, we did not adopt such finer categories here because subdividing social strategies reduces the statistical power of evolutionary models (ref) and focusing only on eusociality would leave too few independent evolutionary origins to evaluate broad evolutionary patterns. In addition, many of the empirical links between climate and sociality in bees come from non-eusocial taxa such as sweat bees (Halictidae) and *Ceratina*, so restricting our analyses to eusocial lineages alone would overlook much of the relevant ecological variation (Davison & Field 2017; Wcislo & Danforth 1997; Eickwort et al. 1996; Groom & Rehan 2018). For example, in Halictidae, social populations are more common in regions with longer warm seasons that support extended foraging periods (Davison & Field 2017), while shifts northward or to higher elevations are often accompanied by transitions from social to solitary nesting (Wcislo & Danforth 1997), sometimes even within the same species (Eickwort et al. 1996). Similarly, in *Ceratina*, social species tend to have tropical distributions (Groom & Rehan 2018). Another potentially informative distinction is between cavity nesting and soil excavation, which could help clarify the relationship between nesting biology and precipitation. However, because cavity nesting is closely associated with above-ground nesting for most species, this signal is largely encompassed by the above-versus below-ground classification, so only limited additional resolution would be expected.

Alternatively, temperature may act not only as a direct constraint, but also as a macroevolutionary proxy for the intensity of biotic interactions (Schemske et al. 2009). Soil microbes (including pathogenic fungi and bacteria) and predatory ants, prominent in tropical ecosystems, are ectothermic organisms whose activity, abundance, and metabolic rates are directly governed by ambient temperature and its stability (Brown et al. 2004). Sociality may help buffer bees against predation by ants through improved nest defense, while nesting above-ground reduces exposure to soil-associated microbes. Therefore, the evolution of the social, above-ground nesting state in tropical bees may also reflect avoidance of the high biotic risks these climates promote rather than direct tracking of temperature per se. From this perspective, temperature captures both the abiotic environment and the ecological context of biotic interactions, which may explain why temperature, rather than precipitation, emerges as the dominant axis of evolutionary divergence in sociality and nesting strategy.

### Evolutionary constraints and niche conservatism shape bimodal latitudinal diversity gradient in bees

Unlike many other taxa that reach their greatest diversity in the tropics (Hillebrand 2004), bees exhibit a bimodal latitudinal diversity gradient, peaking in temperate and arid habitats (Orr et al. 2021). Traditional latitudinal diversity gradients (LDG) have been explained through a variety of mechanisms (Mittelbach et al. 2007), among them tropical niche conservatism, or the out-of-tropics hypothesis (Jablonski et al. 2006; Wiens & Donoghue 2004, 2025). This hypothesis posits that clades tend to be most diverse in the tropics because tropical environments are older and were historically more widespread, leading many clades to originate there. Over time, these clades often remain associated with stable tropical environments because they evolve narrow physiological tolerances that restrict dispersal into temperate regions, giving rise to the traditional LDG observed across many groups (Janzen 1967; Addo-Bediako et al. 2000).

Similarly, this logic can be applied to trait-based richness patterns, in that ancestral trait states tend also to be the most common in a clade (Wiens & Donoghue 2004, 2025). This pattern likely underlies the LDG observed in flowering plants, which originated in the tropics and appear to have retained this ancestral climatic niche across many lineages (Wiens & Donoghue 2025).

These same mechanisms can be invoked to explain exceptions to the traditional LDG, such as the bimodal LDG seen in bees. Fossil and phylogenomic evidence indicate that bees originated in Western Gondwana during the Cretaceous, which at the time was likely characterized by a xeric climate—conditions that also characterize modern hotspots of bee species richness (Almeida et al. 2023; Michener 1979; Orr et al. 2021). Bees are also likely ancestrally solitary and ground-nesting (Sann et al. 2018; Debevec et al. 2012; Branstetter et al. 2017; Michener 1974, 2007), which remains the most speciose trait combination in the present day. These observations suggest that ancestral bees may have been adapted to arid environments, and that most extant bee species have retained these ancestral niche characteristics, a pattern known as phylogenetic niche conservatism (Wiens & Donoghue 2004). In this context, the bimodal LDG in bees may be explained not by the tropics being inherently unsuitable, but by the fact that the ancestral suite of bee traits is poorly adapted for tropical conditions and the derived traits required for tropical success have evolved infrequently.

Indeed, we find that the joint evolution of sociality and above-ground nesting, the strategy that is most prevalent in tropical environments, occurs rarely across the evolutionary history of bees. However, once present, bees with both traits show the narrowest climatic niche breadths of all trait combinations. Although it remains unclear whether the narrow climatic niches of social, above-ground nesting bees are the result of physiological dispersal limitations or historical biogeography, this finding aligns with theory predicting a narrowing of climatic niches at lower latitudes, usually invoked as an explanation for dispersal limitations of tropical taxa and the emergence of traditional LDGs (MacArthur 1984). By contrast, social, above-ground nesting bees appear to have narrowed their climatic niches following colonization of the tropics after originating in arid, temperate habitats, rather than the reverse pattern seen in many other groups. Importantly, for bees this transition seems to have been facilitated by the evolution of key traits allowing them to expand into and persist in tropical regions, highlighting how the evolutionary dynamics of lineage-specific key traits can mediate broader trends in niche breadth and diversity gradients that emerge across taxa and biomes.

Finally, although the solitary, ground-nesting state is ancestral in bees, some social, above-ground lineages diverged early in the evolutionary history of bees. For example, the oldest known bee fossil, *Cretotrigona prisca*, is a stingless bee (Engel 2000) and model-based estimates place the origin of eusociality in the corbiculate bees at least 87 Mya (Cardinal & Danforth, 2011). Therefore, a question that remains to be answered is: why have social, above-ground nesting bee lineages in the tropics not undergone diversification that parallels that of solitary, ground-nesting bee lineages outside the tropics, which are of similar evolutionary age?

In addition to phylogenetic niche conservatism and evolutionary constraints, low tropical bee diversity may also stem from emergent properties of the social, above-ground nesting state that dominates in these regions, specifically the way bees with these traits structure bee communities. Although tropical social bees have narrow climatic niches, they are hyper-generalist foragers, likely filling a broad range of ecological roles in tropical pollination networks that might otherwise be occupied by a diverse assemblage of more specialized solitary species (Roubik 1992; Ogilvie & Forrest 2017). For example, across tropical forests, stingless bees are numerically abundant and competitively dominant, comprising a large proportion (∼50–70%) of bees observed foraging on flowers, sometimes outcompeting other bees (Garibaldi et al. 2021; Cairns et al. 2005; Brosi et al. 2008; Ramalho 2004; Roubik 1992 and citations therein). Therefore, the prevalence of social, above-ground nesting bees in the tropics despite relatively low bee diversity in these regions overall may reflect the ecological dominance of a few highly successful lineages, such as Meliponini (Roubik 1992). These dynamics may be one reason why, despite low tropical bee richness, the proportion of bee-pollinated flowers (∼50%) is comparable to that in regions with much higher bee diversity, perhaps reflecting reliance on fewer abundant, generalist lineages (Martins et al. 2025).

### Future directions

Future research should aim to clarify the mechanisms linking sociality and above-ground nesting to tropical environments, including testing the relative roles of predation pressure, competition with and among highly social lineages, floral resource dynamics, and pollen spoilage in shaping the functional composition of tropical bee communities. Addressing these questions will require integrative approaches that combine natural history observations, experimental manipulations, and comparative phylogenetic analyses, alongside efforts to expand trait and occurrence datasets for understudied groups. For example, targeted studies of clades that are highly labile in these life history traits such as *Lasioglossum* could help clarify the evolutionary relationships between climate and life history traits in bees, yet natural history data for these groups is limited.

Additionally, although macroecological patterns of bee diversity are likely robust to sampling biases (Orr et al. 2021), our understanding of the dynamics governing tropical bee diversity patterns will be improved by exploring sampling gaps in tropical rainforests with methods that better capture vertical stratification in bee communities, as many tropical bees predominantly forage from the canopy where they are difficult to collect (Dorey et al. 2024; Cunningham-Minnick et al. 2024). The use of plant materials such as oils and resins in tropical bee nest construction also merits further study, as these behaviors may facilitate persistence in humid tropical environments where floral oil and resin-collecting bees are most diverse (Michener 2007; Martins et al. 2015; Danforth et al. 2019; Buchmann 1987). Finally, our findings highlight variability in the functional composition of bee communities across space, which may result in divergent selection pressures on the floral morphologies and rewards of bee-pollinated plants, depending on how bee traits influence floral preferences. As our understanding of bee biogeography and traits improve, so too will our ability to understand how geographic variation in bee communities shapes pollination dynamics across ecosystems.

### Conclusions

We find that social, above-ground nesting bees are closely tied to tropical environments, yet the evolution of these derived traits is highly constrained, unfolding only through rare, stepwise transitions from the ancestral solitary, ground-nesting state. Unlike many groups where tropical richness accumulates through limited dispersal out of the tropics, bees appear to have retained ancestral traits favoring arid and seasonal habitats, with only rare transitions enabling persistence in the tropics. When these transitions did occur, they produced highly abundant but relatively species-poor assemblages dominated by social, above-ground nesting bees. Together, these findings suggest that bee diversity patterns are shaped by the interaction of life history traits, climate, and evolutionary history, providing insight into why bees deviate from the canonical latitudinal diversity gradient and, more broadly, how clade-specific evolutionary trajectories and the ecological consequences of key traits can generate exceptions to this otherwise pervasive biodiversity pattern.

## Acknowledgements

We thank the Vasconcelos lab, James Boyko, the floral specialization working group, and L. Heinrich’s dissertation committee members (Nate Sanders, Marjorie Weber, and Brian Weeks) for their thoughtful feedback throughout this project.

## References

Addo-Bediako, A., S. L. Chown, and K. J. Gaston. (2000). Thermal tolerance, climatic variability and latitude. Proceedings of the Royal Society of London, Series B: Biological Sciences 267: 739–745.

Aguiar, A. J. C., G. A. R. Melo, T. N. C. Vasconcelos, R. B. Gonçalves, L. Giugliano, and A. C. Martins. (2020). Biogeography and early diversification of Tapinotaspidini oil-bees support presence of Paleocene savannas in South America. Molecular Phylogenetics and Evolution 143: 106692. 10.1016/j.ympev.2019.106692

Almeida, E. A. B., S. Bossert, B. N. Danforth, D. S. Porto, F. V. Freitas, C. C. Davis, E. A. Murray, B. B. Blaimer, T. Spasojevic, P. R. Ströher, M. C. Orr, L. Packer, S. G. Brady, M. Kuhlmann, M. G. Branstetter, and M. R. Pie. (2023). The evolutionary history of bees in time and space. Current Biology 33(16): 3409–3422.e6. 10.1016/j.cub.2023.07.005

Antoine, C. M., and J. R. K. Forrest. (2021). Nesting habitat of ground-nesting bees: a review. Ecological Entomology 46: 143–159.

Beaulieu, J. M., B. C. O’Meara, and M. J. Donoghue. (2013). Identifying hidden rate changes in the evolution of a binary morphological character: the evolution of plant habit in campanulid angiosperms. Systematic Biology 62:725–737. 10.1093/sysbio/syt034

Bossert, S., T. J. Wood, S. Patiny, D. Michez, E. A. B. Almeida, R. L. Minckley, L. Packer, J. L. Neff, R. S. Copeland, J. Straka, A. Pauly, T. Griswold, S. G. Brady, B. N. Danforth, and E. A. Murray. (2022). Phylogeny, biogeography and diversification of the mining bee family Andrenidae. Systematic Entomology 47: 283–302.

Boyko, J. D. (2024). Automatic discovery of optimal discrete character models. bioRxiv. 10.1101/2024.11.15.623760

Boyko, J. D., & Beaulieu, J. M. (2023). Reducing the biases in false correlations between discrete characters. Systematic Biology 72(2), 476–488.

Boyko, J. D., Hagen, E. R., Beaulieu, J. M., & Vasconcelos, T. (2023). The evolutionary responses of life-history strategies to climatic variability in flowering plants. New Phytologist 240(4): 1587–1600.

Boyko, J. D., B. C. O’Meara, and J. M. Beaulieu. (2023). A novel method for jointly modeling the evolution of discrete and continuous traits. Evolution 77: 836–851.

Branstetter, M. G., B. N. Danforth, J. P. Pitts, B. C. Faircloth, P. S. Ward, M. L. Buffington, M. W. Gates, R. R. Kula, and S. G. Brady. (2017). Phylogenomic insights into the evolution of stinging wasps and the origins of ants and bees. Current Biology 27: 1019–1025.

Brosi, B. J., G. C. Daily, T. M. Shih, F. Oviedo, and G. Durán. (2008). The effects of forest fragmentation on bee communities in tropical countryside. Journal of Applied Ecology 45(3): 773–783. 10.1111/j.1365-2664.2007.01412.x

Brown, J. H., J. F. Gillooly, A. P. Allen, V. M. Savage, and G. B. West. (2004). Toward a metabolic theory of ecology. Ecology 85: 1771–1789.

Buchmann, S. L. (1987). The ecology of oil flowers and their bees. Annual Review of Ecology and Systematics 18:343–369.

Bueno, F. G. B., L. Kendall, D. A. Alves, M. L. Tamara, T. Heard, T. Latty, and R. Gloag. (2023). Stingless bee floral visitation in the global tropics and subtropics. Global Ecology and Conservation 43: e02454. 10.1016/j.gecco.2023.e02454

Cairns, C. E., R. Villanueva-Gutiérrez, S. Koptur, and D. B. Bray. (2005). Bee populations, forest disturbance, and africanization in Mexico. Biotropica 37(4): 686–692. 10.1111/j.1744-7429.2005.00087.x

Cane, J. H., and J. L. Neff. (2011). Predicted fates of ground-nesting bees in soil heated by wildfire: Thermal tolerances of life stages and a survey of nesting depths. Biological Conservation 144: 2631–2636. 10.1016/j.biocon.2011.07.019

Cardinal, S., and B. N. Danforth. (2011). The antiquity and evolutionary history of social behavior in bees. PLoS ONE 6(6): e21086. 10.1371/journal.pone.0021086

Cardinal, S., and B. N. Danforth. (2013). Bees diversified in the age of eudicots. Proceedings of the Royal Society B: Biological Sciences 280(1755): 20122686. 10.1098/rspb.2012.2686

Cheng, W., and L. Ashton. (2021). Ecology: what affects the distribution of global bee diversity. Current Biology 31: R127–R128.

Chui, S. X., A. Keller, and S. D. Leonhardt. (2022). Functional resin use in solitary bees. Ecological Entomology 47(2): 115–136. 10.1111/een.13103

Cunningham-Minnick, M. J., H. P. Roberts, J. Milam, and D. I. King. (2024). Sampling the understory, midstory, and canopy is necessary to fully characterize native bee communities of temperate forests and their dynamic environmental relationships. Frontiers in Ecology and Evolution 12. 10.3389/fevo.2024.1352266

Danforth, B. N., S. Cardinal, C. Praz, E. A. B. Almeida, and D. Michez. (2013). The impact of molecular data on our understanding of bee phylogeny and evolution. Annual Review of Entomology 58: 57–78.

Danforth, B. N., R. L. Minckley, and J. L. Neff. (2019). The solitary bees: biology, evolution, conservation. Princeton University Press, Princeton, New Jersey, USA.

da Silva, J. (2021). Life history and the transitions to eusociality in the Hymenoptera. Frontiers in Ecology and Evolution 9.

Davison, P. J., and J. Field. (2017). Season length, body size, and social polymorphism: size clines but not saw tooth clines in sweat bees. Ecological Entomology 42: 768–776.

Debevec, A. H., S. Cardinal, and B. N. Danforth. (2012). Identifying the sister group to the bees: a molecular phylogeny of Aculeata with an emphasis on the superfamily Apoidea. Zoologica Scripta 41:527–535.

Devkota, K., C. F. dos Santos, P. D. Souza-Santos, J. D. Ramos, A. Otesbelgue, B. P. Mishra, E. A. B. Almeida, and B. Blochtein. (2024). Pollen diet diversity across bee lineages varies with lifestyle rather than colony size. Journal of Insect Science 24(2): 1. 10.1093/jisesa/ieae023

Dorey, J. B., E. E. Fischer, P. R. Chesshire, A. Nava-Bolaños, R. L. O’Reilly, S. Bossert, S. M. Collins, E. M. Lichtenberg, E. M. Tucker, A. Smith-Pardo, A. Falcon-Brindis, D. A. Guevara, B. Ribeiro, D. de Pedro, J. Pickering, K.-L. J. Hung, K. A. Parys, L. M. McCabe, M. S. Rogan, R. L. Minckley, S. J. E. Velazco, T. Griswold, T. A. Zarrillo, W. Jetz, Y. V. Sica, M. C. Orr, L. M. Guzman, J. S. Ascher, A. C. Hughes, and N. S. Cobb. (2023). A globally synthesised and flagged bee occurrence dataset and cleaning workflow. Scientific Data 10: 747.

Dorey, J., O. Davies, K. Magnacca, M. Schwarz, A.-M. Gilpin, T. Ramage, M. Tuiwawa, S. Groom, M. Stevens, and B. Parslow. (2024). Canopy specialist Hylaeus bees highlight sampling biases and resolve Michener’s mystery. Frontiers in Ecology and Evolution 12. 10.3389/fevo.2024.1339446

Eickwort, G. C., J. M. Eickwort, J. Gordon, M. A. Eickwort, and W. T. Wcislo. (1996). Solitary behavior in a high-altitude population of the social sweat bee *Halictus rubicundus* (Hymenoptera: Halictidae). Behavioral Ecology and Sociobiology 38(4): 227–233. 10.1007/s002650050236

Engel, M. S. (2000). A new interpretation of the oldest fossil bee (Hymenoptera, Apidae). American Museum Novitates 3296: 1–11.

Evans, H. E. (1977). Commentary: Extrinsic versus intrinsic factors in the evolution of insect sociality. BioScience 27(9): 613–617. 10.2307/1297657

Felsenstein, J. (1985). Phylogenies and the comparative method. The American Naturalist 125: 1–15.

Freitas, F. V., M. G. Branstetter, T. Griswold, and E. A. B. Almeida. (2021). Partitioned gene-tree analyses and gene-based topology testing help resolve incongruence in a phylogenomic study of host-specialist bees (Apidae: Eucerinae). Molecular Biology and Evolution 38: 1090–1100.

Garibaldi, L. A., N. Pérez-Méndez, G. D. Cordeiro, A. Hughes, M. Orr, I. Alves-Dos-Santos, B. M. Freitas, F. Freitas de Oliveira, G. LeBuhn, I. Bartomeus, M. A. Aizen, P. B. Andrade, B. Blochtein, D. Boscolo, P. M. Drumond, M. C. Gaglianone, B. Gemmill-Herren, R. Halinski, C. Krug, M. M. Maués, L. H. P. Kiill, M. Pinheiro, C. S. S. Pires, and B. F. Viana. (2021). Negative impacts of dominance on bee communities: Does the influence of invasive honey bees differ from native bees? Ecology 102(12): e03526. 10.1002/ecy.3526

Goulson, D. (2003). Effects of introduced bees on native ecosystems. Annual Review of Ecology, Evolution, and Systematics 34: 1–26.

Groom, S., and S. Rehan. 2018. Climate-mediated behavioural variability in facultatively social bees. Biological Journal of the Linnean Society 125: 165–170.

Gruchowski-Woitowicz, F. C., C. I. da Silva, and M. Ramalho. (2024). Influence of generalist stingless bees on the structure of mutualistic flower–pollinator networks in the tropics: Temporal variation. Ecological Entomology 49(3): 338–356. 10.1111/een.13308

Grüter, C., and L. Hayes. (2022). Sociality is a key driver of foraging ranges in bees. Current Biology 32(24): 5390–5397.e3. 10.1016/j.cub.2022.10.064

Grüter, C. (2020). Stingless Bees: Their Behaviour, Ecology and Evolution. Springer International Publishing, Cham.

Grüter, C., F. H. I. D. Segers, C. Menezes, A. Vollet-Neto, T. Falcón, L. von Zuben, M. M. G. Bitondi, F. S. Nascimento, and E. A. B. Almeida. (2017). Repeated evolution of soldier sub-castes suggests parasitism drives social complexity in stingless bees. Nature Communications 8: 4.

Heinrich, B. (1972). Temperature regulation in the bumblebee *Bombus vagans*: A field study. Science 175: 185–187.

Heinrich, B. (1975). Thermoregulation in bumblebees. Journal of Comparative Physiology 96: 155–166.

Henríquez-Piskulich, P., A. F. Hugall, and D. Stuart-Fox. (2024). A supermatrix phylogeny of the world’s bees (Hymenoptera: Anthophila). Molecular Phylogenetics and Evolution 190: 107963. 10.1016/j.ympev.2023.107963

Hijmans, R. J. (2022). *raster: Geographic data analysis and modeling*. R package version 3.5-13. Retrieved from https://CRAN.R-project.org/package=raster

Hillebrand, H. (2004). On the generality of the latitudinal diversity gradient. The American Naturalist 163: 192–211.

Hines, H.M. (2008) Historical biogeography, divergence times, and diversification patterns of bumble bees (Hymenoptera: Apidae: Bombus). Systematic Biology 57: 58–75

Jablonski, D., K. Roy, and J. W. Valentine. (2006). Out of the tropics: Evolutionary dynamics of the latitudinal diversity gradient. Science 314: 102–106.

Janzen, D. H. (1967). Why mountain passes are higher in the tropics. The American Naturalist 101:233–249.

Jeanne, R. L. (1979). A latitudinal gradient in rates of ant predation. Ecology 60: 1211–1224.

Kapheim, K. M., A. R. Smith, P. Nonacs, W. T. Wcislo, and R. K. Wayne. (2013). Foundress polyphenism and the origins of eusociality in a facultatively eusocial sweat bee, Megalopta genalis (Halictidae). Behavioral Ecology and Sociobiology 67: 331–340.

Karger, D. N., O. Conrad, J. Böhner, T. Kawohl, H. Kreft, R. W. Soria-Auza, N. E. Zimmermann, H. P. Linder, and M. Kessler. (2017). Climatologies at high resolution for the earth’s land surface areas. Scientific Data 4(1): 170122. 10.1038/sdata.2017.122

King, B., and M. S. Y. Lee. (2015). Ancestral state reconstruction, rate heterogeneity, and the evolution of reptile viviparity. Systematic Biology 64:532–544. 10.1093/sysbio/syv005

Kocher, S. D., L. Pellissier, C. Veller, J. Purcell, M. A. Nowak, M. Chapuisat, and N. E. Pierce. (2014). Transitions in social complexity along elevational gradients reveal a combined impact of season length and development time on social evolution. Proceedings of the Royal Society B: Biological Sciences 281: 20140627. 10.1098/rspb.2014.0627

Krombein, K. V., P. D. Hurd, D. R. Smith, and B. D. Burks. 1979. Catalog of Hymenoptera in America North of Mexico. Volume 2. Smithsonian Institution Press, Washington, D.C., USA.

Lafarge, T., and B. Pateiro-Lopez. 2023. alphashape3d: Implementation of the 3D Alpha-Shape for the Reconstruction of 3D Sets from a Point Cloud. R package version 1.3–2. 10.32614/CRAN.package.alphashape3d

Lin, N., and C. D. Michener. (1972). Evolution of sociality in insects. The Quarterly Review of Biology 47(2): 131–159. 10.1086/407216

Lokvam, J., and J. F. Braddock. (1999). Anti-bacterial function in the sexually dimorphic pollinator rewards of *Clusia grandiflora* (Clusiaceae). Oecologia 119: 534–540. 10.1007/s004420050885

MacArthur, R. H. 1984. Geographical ecology: Patterns in the distribution of species. Princeton University Press, Princeton, New Jersey, USA.

Martins, A. C., G. A. R. Melo, and S. S. Renner. (2015). Gain and loss of specialization in two oil-bee lineages, *Centris* and *Epicharis* (Apidae). Evolution 69(7): 1835–1844. 10.1111/evo.12689

Martins, A. C., C. E. B. Proença, T. N. C. Vasconcelos, A. J. C. Aguiar, H. C. Farinasso, A. T. F. de Lima, J. E. Q. Faria, K. Norrana, M. B. R. Costa, M. M. Carvalho, R. L. Dias, M. M. C. Bustamante, F. A. Carvalho, and A. Keller. (2023). Contrasting patterns of foraging behavior in neotropical stingless bees using pollen and honey metabarcoding. Scientific Reports 13(1): 14474. 10.1038/s41598-023-41304-0

Martins, A. C., L. Heinrich, A. C. Hughes, K. C. Seltmann, M. C. Orr, and T. Vasconcelos. (2025). Plant communities in the Americas are highly bee dependent regardless of biome or local bee diversity. Global Ecology and Biogeography 34: e70101.

Michener, C. D. (1969). Comparative social behavior of bees. In Annual Review of Entomology 14: 299–342. Annual Reviews. 10.1146/annurev.en.14.010169.001503

Michener, C. D. (1974). The social behavior of the bees: A comparative study. Belknap Press of Harvard University Press, Cambridge, Massachusetts.

Michener, C. D. (1979). Biogeography of the bees. Annals of the Missouri Botanical Garden 66(3): 277–347. 10.2307/2398833

Michener, C. D. (2007). The Bees of the World, 2nd ed. Johns Hopkins University Press, Baltimore, Maryland.

Mikát, M., K. Černá, and J. Straka. (2016). Major benefits of guarding behavior in subsocial bees: Implications for social evolution. Ecology and Evolution 6: 6784–6797. 10.1002/ece3.2463

Minckley, R. L., J. H. Cane, and L. Kervin. (2000). Origins and ecological consequences of pollen specialization among desert bees. Proceedings of the Royal Society B: Biological Sciences 267: 265–271.

Mittelbach, G. G., D. W. Schemske, H. V. Cornell, A. P. Allen, J. M. Brown, M. B. Bush, S. P. Harrison, A. H. Hurlbert, N. Knowlton, H. A. Lessios, C. M. McCain, A. R. McCune, L. A. McDade, M. A. McPeek, T. J. Near, T. D. Price, R. E. Ricklefs, K. Roy, D. F. Sax, D. Schluter, J. M. Sobel, and M. Turelli. (2007). Evolution and the latitudinal diversity gradient: speciation, extinction and biogeography. Ecology Letters 10: 315–331.

Norden, B. B., K. V. Krombein, M. A. Deyrup, and J. P. Edirisinghe. (2003). Biology and behavior of a seasonally aquatic bee, Perdita (Alloperdita) floridensis (Hymenoptera: Andrenidae: Panurginae). Journal of the Kansas Entomological Society 76: 236–249.

Ogilvie, J. E., and J. R. Forrest. (2017). Interactions between bee foraging and floral resource phenology shape bee populations and communities. Current Opinion in Insect Science 21: 75–82.

Oksanen, J., G. Simpson, F. Blanchet, R. Kindt, P. Legendre, P. Minchin, R. O’Hara, P. Solymos, M. Stevens, E. Szoecs, H. Wagner, M. Barbour, M. Bedward, B. Bolker, D. Borcard, G. Carvalho, M. Chirico, M. De Caceres, S. Durand, H. Evangelista, R. FitzJohn, M. Friendly, B. Furneaux, G. Hannigan, M. Hill, L. Lahti, D. McGlinn, M. Ouellette, E. Ribeiro Cunha, T. Smith, A. Stier, C. Ter Braak, J. Weedon, and T. Borman. 2025. vegan: Community Ecology Package. R package version 2.6–10. 10.32614/CRAN.package.vegan

Orr, M. C., A. C. Hughes, D. Chesters, J. Pickering, C.-D. Zhu, and J. S. Ascher. (2021). Global patterns and drivers of bee distribution. Current Biology 31: 451–458.e4. 10.1016/j.cub.2020.11.056

Ostwald, M. M., V. H. Gonzalez, C. Chang, N. Vitale, M. Lucia, and K. C. Seltmann. (2024). Toward a functional trait approach to bee ecology. Ecology and Evolution 14: e70465.

Pagel, M. (1994). Detecting correlated evolution on phylogenies: a general method for the comparative analysis of discrete characters. Proceedings of the Royal Society of London. Series B: Biological Sciences 255(1342): 37–45.

Pebesma, E., and R. S. Bivand. (2005). sp: Classes and methods for spatial data. R package version 1.4-5. Retrieved from https://CRAN.R-project.org/package=sp

Peled, O., G. Greenbaum, and G. Bloch. (2024). Data-driven analyses of social complexity in bees reveal phenotypic diversification following a major evolutionary transition. Preprint, Evolutionary Biology.

Potts, S., and P. Willmer. (1997). Abiotic and biotic factors influencing nest-site selection by *Halictus rubicundus*, a ground-nesting halictine bee. Ecological Entomology 22(3): 319–328. 10.1046/j.1365-2311.1997.00071.x

Purcell, J. (2011). Geographic patterns in the distribution of social systems in terrestrial arthropods. Biological Reviews 86: 475–491.

Ramalho, M. (2004). Stingless bees and mass flowering trees in the canopy of Atlantic Forest: A tight relationship. Acta Botanica Brasilica 18: 37–47. 10.1590/S0102-33062004000100005

Roubik, D. W. (1989) Ecology and natural history of tropical bees. Cambridge University Press, New York.

Roubik, D. W. (2023) Stingless bee (Apidae: Apinae: Meliponini) ecology. Annual Review of Entomology 68: 231–256.

Roubik, D.W. & Michener, C. (1980). The seasonal cycle and nests of *Epicharis zonata*, a bee whose cells are below the wet-Season water table (Hymenoptera: Anthophoridae). Biotropica. 12(1): 56–60. 10.2307/2387773

Roubik, D. W. (1992). Loose niches in tropical communities: Why are there so few bees and so many trees? In M. D. Hunter, T. Ohgushi, and P. W. Price (Eds.), Effects of Resource Distribution on Animal–Plant Interactions (pp. 327–354). Academic Press. 10.1016/B978-0-08-091881-5.50014-1

Russell, A. L., S. L. Buchmann, J. S. Ascher, Z. Wang, R. Kriebel, D. D. Jolles, M. C. Orr, and A. C. Hughes. (2024). Global patterns and drivers of buzzing bees and poricidal plants. Current Biology 34: 3055–3063.e5.

Sann, M., O. Niehuis, R. S. Peters, C. Mayer, A. Kozlov, L. Podsiadlowski, S. Bank, K. Meusemann, B. Misof, C. Bleidorn, and M. Ohl. (2018). Phylogenomic analysis of Apoidea sheds new light on the sister group of bees. BMC evolutionary biology 18: 71.

Schemske, D. W., G. G. Mittelbach, H. V. Cornell, J. M. Sobel, and K. Roy. (2009). Is there a latitudinal gradient in the importance of biotic interactions? Annual Review of Ecology, Evolution, and Systematics 40: 245–269.

Sedivy, C., S. Dorn, and A. Müller. (2013). Molecular phylogeny of the bee genus *Hoplitis* (Megachilidae: Osmiini)—How does nesting biology affect biogeography? Zoological Journal of the Linnean Society 167(1): 28–42. 10.1111/j.1096-3642.2012.00876.x

Sless, T. J. L., M. G. Branstetter, J. P. Gillung, E. A. Krichilsky, K. B. Tobin, J. Straka, J. G. Rozen, F. V. Freitas, A. C. Martins, S. Bossert, J. B. Searle, and B. N. Danforth. (2022). Phylogenetic relationships and the evolution of host preferences in the largest clade of brood parasitic bees (Apidae: Nomadinae). Molecular Phylogenetics and Evolution 166: 107326.

Smith, A. R., W. T. Wcislo, and S. O’Donnell. (2003). Assured fitness returns favor sociality in a mass-provisioning sweat bee, *Megalopta genalis* (Hymenoptera: Halictidae). Behavioral Ecology and Sociobiology 54: 14–21. 10.1007/s00265-003-0650-2

Spessa, A., M. P. Schwarz, and M. Adams. (2000). Sociality in *Amphylaeus morosus* (Hymenoptera: Colletidae: Hylaeinae). Annals of the Entomological Society of America 93(3): 684–692. 10.1603/0013-8746(2000)093[0684:SIAMHC]2.0.CO;2

Staab, M., Pufal, G., Tscharntke, T., & Klein, A. M. (2018). Trap nests for bees and wasps to analyse trophic interactions in changing environments—A systematic overview and user guide. Methods in Ecology and Evolution, 9(11): 2226–2239.

TDWG World Geographic Scheme for Recording Plant Distributions Committee. (2001). World Geographic Scheme for Recording Plant Distributions Standard. Biodiversity Information Standards (TDWG). Retrieved from http://www.tdwg.org/standards/109

Vasconcelos, T., Boyko, J. D., & Beaulieu, J. M. (2023). Linking mode of seed dispersal and climatic niche evolution in flowering plants. Journal of Biogeography, 50(1), 43–56.

Vogel, S. (1974). Ölblumen und ölsammelnde Bienen. Tropische und subtropische Pflanzenwelt. Akademie der Wissenschaften und der Literatur, Franz Steiner Verlag, Wiesbaden.

Wcislo, W. T., and B. N. Danforth. (1997). Secondarily solitary: The evolutionary loss of social behavior. Trends in Ecology & Evolution 12(12): 468–474. 10.1016/S0169-5347(97)01198-1

Wcislo, W.T. (1987). The roles of seasonality, host synchrony, and behavior in the evolutions and distributions of nest parasites in Hymenoptera (Insecta), with special reference to bees (Apoidea). Biological Reviews, 62: 515–542. 10.1111/j.1469-185X.1987.tb01640.x

Wiens, J. J., and M. J. Donoghue. (2004). Historical biogeography, ecology and species richness. Trends in Ecology & Evolution 19: 639–644. 10.1016/j.tree.2004.09.011

Wiens, J. J., and M. J. Donoghue. (2025). The origins of the latitudinal diversity gradient: Revisiting the tropical conservatism hypothesis. Journal of Biogeography e15172. 10.1111/jbi.15172

Wiens, J. J. (2023). Trait-based species richness: ecology and macroevolution. Biological Reviews 98:1365–1387.

Williams, P. H. (1998) An annotated checklist of bumble bees with an analysis of patterns of description (Hymenoptera: Apidae, Bombini). Bulletin of Natural History Museum, Entomological Series 67: 79–152

