## Supplemental Materials for "Sociality and nesting strategy shape the bimodal diversity gradient in bees"

**Article title:** Sociality and nesting strategy shape the bimodal diversity gradient in bee distribution

**Table S1 (separate PDF).** List of species with known history of introductions outside their native range which were excluded from analyses.

**Table S2 (separate Excel spreadsheet).** Trait dataset used in the study, including nesting biology and sociality data for 4,293 bee species sampled in the Henríquez-Piskulich et al. (2024) phylogenetic tree, along with literature references for each scoring.

**Table S3 (separate PDF).** Kruskal-Wallis post-hoc contrasts.

**Table S4.** Summary of average transition counts and ages based on results from corHMM ancestral state reconstructions. Rows are ordered from the rarest to the most common transitions, according to the mean number of transitions per stochastic map.

| From | To | Avg. Count | SD Count | Avg. Age | SD Age |
|---|---|---|---|---|---|
| R2 social—ground | R2 social—aboveground | 1.00 | 0.00 | 1.81 | 1.00 |
| R1 solitary—aboveground | R1 social—aboveground | 1.71 | 0.83 | 40.92 | 30.17 |
| R2 solitary—ground | R2 solitary—aboveground | 3.44 | 1.09 | 16.74 | 13.68 |
| R2 social—aboveground | R2 solitary—aboveground | 8.10 | 1.87 | 5.70 | 6.93 |
| R1 solitary—ground | R1 social—ground | 9.61 | 1.76 | 16.32 | 12.12 |
| R2 solitary—aboveground | R2 social—aboveground | 12.00 | 1.60 | 11.79 | 18.03 |
| R1 social—aboveground | R1 social—ground | 15.31 | 0.56 | 7.75 | 10.84 |
| R1 solitary—ground | R1 solitary—aboveground | 19.51 | 3.79 | 26.26 | 27.06 |
| R1 social—ground | R2 social—ground | 19.96 | 4.09 | 7.38 | 8.01 |
| R1 social—ground | R1 solitary—ground | 24.05 | 2.14 | 5.82 | 6.91 |
| R1 solitary—aboveground | R2 solitary—aboveground | 48.13 | 7.42 | 20.99 | 21.58 |
| R2 solitary—aboveground | R1 solitary—aboveground | 52.50 | 6.09 | 11.90 | 11.94 |
| R1 solitary—aboveground | R1 solitary—ground | 72.27 | 6.73 | 19.47 | 22.88 |
| R1 solitary—ground | R2 solitary—ground | 104.98 | 12.26 | 25.29 | 23.76 |

**Table S5.** Model comparison results for each climate variable. The best-fitting model for each variable (lowest AICc) is bolded. Models are sorted by AICc within each variable.

| Variable | Model (Description) | np | lnLik | DiscLik | ContLik | AICc | dAICc | AICcwt |
|---|---|---|---|---|---|---|---|---|
| **BIO15 — Precipitation seasonality** | | | | | | | | |
| | **OU: single optimum (no trait structure)** | **13** | **-3181.11** | **-644.70** | **-2531.80** | **6388.31** | **0.00** | **0.9987** |
| | OU: separate optima for trait combinations | 14 | -3186.74 | -647.25 | -2534.83 | 6401.60 | 13.29 | 0.0013 |
| | OU: separate optima for each corHMM-inferred state | 16 | -3192.49 | -637.91 | -2550.01 | 6417.12 | 28.81 | 5.54E-07 |
| | Brownian motion (1 rate) | 12 | -3813.35 | -630.75 | -3177.99 | 7650.78 | 1262.47 | 7.22E-275 |
| **BIO12 — Annual precipitation** | | | | | | | | |
| | **OU: separate optima for trait combinations** | **16** | **-4247.30** | **-612.94** | **-3629.73** | **8526.74** | **0.00** | **0.9948** |
| | OU: separate optima by nesting strategy | 16 | -4253.14 | -614.45 | -3632.97 | 8538.43 | 11.70 | 0.0029 |
| | OU: single optimum (no trait structure) | 15 | -4254.38 | -609.62 | -3640.16 | 8538.89 | 12.16 | 0.0023 |
| | OU: separate optima for each corHMM-inferred state | 18 | -4255.24 | -615.04 | -3635.66 | 8546.66 | 19.93 | 4.69E-05 |
| | OU: separate optima by sociality | 16 | -4265.83 | -623.74 | -3631.24 | 8563.81 | 37.08 | 8.85E-09 |
| | Brownian motion (1 rate) | 14 | -4600.21 | -611.66 | -3983.94 | 9228.53 | 701.79 | 4.03E-153 |
| **BIO4 — Temperature seasonality** | | | | | | | | |
| | **OU: separate optima for each corHMM-inferred state** | **16** | **-3286.89** | **-660.76** | **-2621.43** | **6605.92** | **0.00** | **1.0000** |
| | OU: single optimum (no trait structure) | 13 | -3314.32 | -657.82 | -2651.90 | 6654.73 | 48.81 | 2.51E-11 |
| | OU: separate optima by sociality | 14 | -3307.30 | -666.65 | -2634.22 | 6642.72 | 36.80 | 1.02E-08 |
| | OU: separate optima by nesting strategy | 14 | -3317.57 | -662.89 | -2649.84 | 6663.25 | 57.33 | 3.56E-13 |
| | OU: separate optima for trait combinations | 14 | -3325.22 | -663.43 | -2657.40 | 6678.55 | 72.63 | 1.69E-16 |
| | Brownian motion (1 rate) | 12 | -3665.86 | -649.46 | -3011.80 | 7355.81 | 749.89 | 1.46E-163 |
| **BIO1 — Mean annual temperature** | | | | | | | | |
| | **OU: separate optima for each corHMM-inferred state** | **18** | **9800.09** | **-621.36** | **10425.87** | **-19564.00** | **0.00** | **0.9759** |
| | OU: separate optima for trait combinations | 16 | 9793.12 | -615.82 | 10413.45 | -19554.10 | 9.90 | 0.0069 |
| | OU: single optimum (no trait structure) | 15 | 9789.88 | -618.37 | 10412.86 | -19549.63 | 14.37 | 0.0007 |
| | OU: separate optima by sociality | 16 | 9793.99 | -611.13 | 10409.73 | -19555.83 | 8.16 | 0.0165 |
| | OU: separate optima by nesting strategy | 16 | 9784.43 | -619.89 | 10408.68 | -19536.72 | 27.28 | 1.16E-06 |
| | Brownian motion (1 rate) | 14 | 9414.51 | -613.27 | 10032.39 | -18800.91 | 763.09 | 1.94E-166 |

**Table S6.** Expected climatic optima estimated using hidden Markov models of correlated trait and climatic niche evolution (hOUwie) for each combination of bee sociality and nesting strategy.

| Trait Combination | Mean annual temp. (°C) | Temp. seasonality (SD × 100) | Annual precip. (mm) |
|---|---|---|---|
| Social / Above-ground | 22.1 | 186 | 665 |
| Social / Ground | 13.8 | 618 | 727 |
| Solitary / Above-ground | 16.3 | 351 | 729 |
| Solitary / Ground | 16.9 | 430 | 500 |

**Table S7.** Pairwise PERMANOVA results comparing univariate climatic niche space among bee life history strategies for **(A)** precipitation niche breadth and **(B)** temperature niche breadth. Niche breadths were calculated by taking the difference between temperate and precipitation extremes (i.e., BIO5 and BIO6 for temperature, and BIO16 and BIO17 for precipitation). *p*-values are based on 999 permutations.

**(A) Precipitation niche breadth (BIO13–BIO14)**

| Group 1 | Group 2 | $R^2$ | F-value | p-value |
|---|---|---|---|---|
| Solitary/Ground | Solitary/Above-ground | 0.05095 | 150.104 | 0.001 |
| Solitary/Ground | Social/Above-ground | 0.23208 | 654.288 | 0.001 |
| Solitary/Ground | Social/Ground | 0.00925 | 23.361 | 0.001 |
| Solitary/Above-ground | Social/Above-ground | 0.09593 | 128.073 | 0.001 |
| Solitary/Above-ground | Social/Ground | 0.01299 | 20.339 | 0.001 |
| Social/Above-ground | Social/Ground | 0.19597 | 222.775 | 0.001 |

**(B) Temperature niche breadth (BIO5–BIO6)**

| Group 1 | Group 2 | $R^2$ | F-value | p-value |
|---|---|---|---|---|
| Solitary/Ground | Solitary/Above-ground | 0.06622 | 198.271 | 0.001 |
| Solitary/Ground | Social/Above-ground | 0.27408 | 817.433 | 0.001 |
| Solitary/Ground | Social/Ground | 0.00020 | 0.511 | 0.504 |
| Solitary/Above-ground | Social/Above-ground | 0.13117 | 182.229 | 0.001 |
| Solitary/Above-ground | Social/Ground | 0.06095 | 100.273 | 0.001 |
| Social/Above-ground | Social/Ground | 0.36445 | 524.120 | 0.001 |

**Table S8.** Multivariate climatic niche volumes for each bee life history strategy, estimated using the 3D convex hull of species positions in principal component space (PC1–PC3). Volumes are reported in arbitrary units corresponding to the volume of climatic space occupied by each group.

| Trait Combination | Niche Volume |
|---|---|
| Solitary / Ground | 242.36 |
| Solitary / Above-ground | 254.04 |
| Social / Ground | 249.97 |
| Social / Above-ground | 68.08 |

**Table S9.** Pairwise PERMANOVA results comparing multivariate climatic niche space among bee life history strategies. Climatic niche space was defined by the first three principal components of 12 bioclimatic variables. Each comparison tests whether two trait groups (combinations of sociality and nesting strategy) occupy significantly different regions of climate space. *p*-values are based on 999 permutations. In the full model, life history strategy explained 11.5% of the total variation in climatic niche space ($R^2 = 0.115$, $p = 0.001$).

| Group 1 | Group 2 | $R^2$ | F-value | p-value |
|---|---|---|---|---|
| Solitary/Ground | Solitary/Above-ground | 0.032 | 92.782 | 0.001 |
| Solitary/Ground | Social/Above-ground | 0.165 | 426.764 | 0.001 |
| Solitary/Ground | Social/Ground | 0.031 | 80.254 | 0.001 |
| Solitary/Above-ground | Social/Above-ground | 0.102 | 137.163 | 0.001 |
| Solitary/Above-ground | Social/Ground | 0.031 | 50.136 | 0.001 |
| Social/Above-ground | Social/Ground | 0.255 | 313.468 | 0.001 |

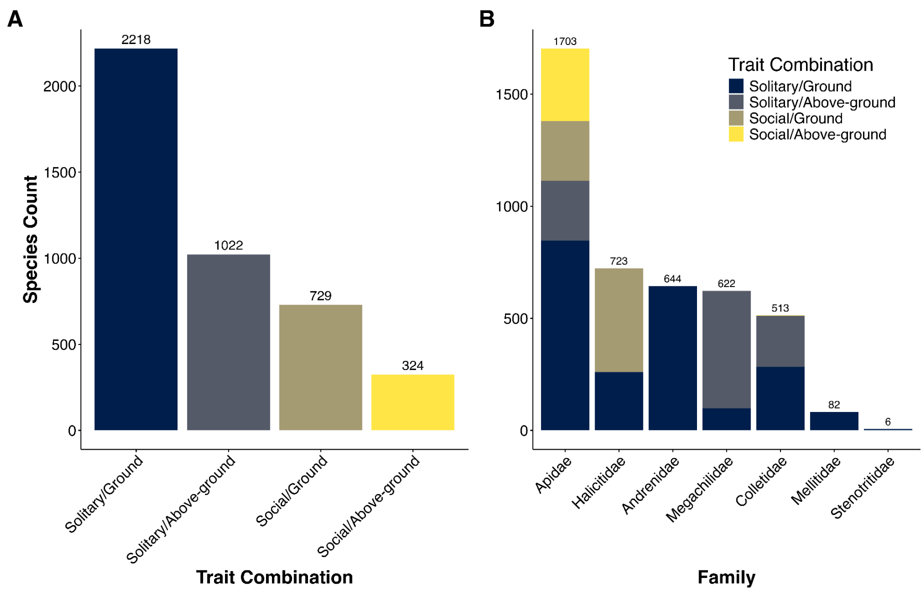

**Figure S1.** Counts of bee species within our trait dataset by **(A)** trait combination (solitary/social and ground/above-ground nesting) and **(B)** family.

**Figure S2 (separate PDF).** Distribution of bees with different sociality/nesting biology combinations across all CHELSA climate variables using global bee occurrence data (Karger et al. 2017). $p < 0.0001$ for all comparisons using Kruskal-Wallis tests.

**Figure S3:** Summary of transition counts for 100 stochastic maps inferred from the corHMMdredge analysis. Note that some of the summarized transitions are between rate classes 1 ("fast") and rate class 2 ("slow").

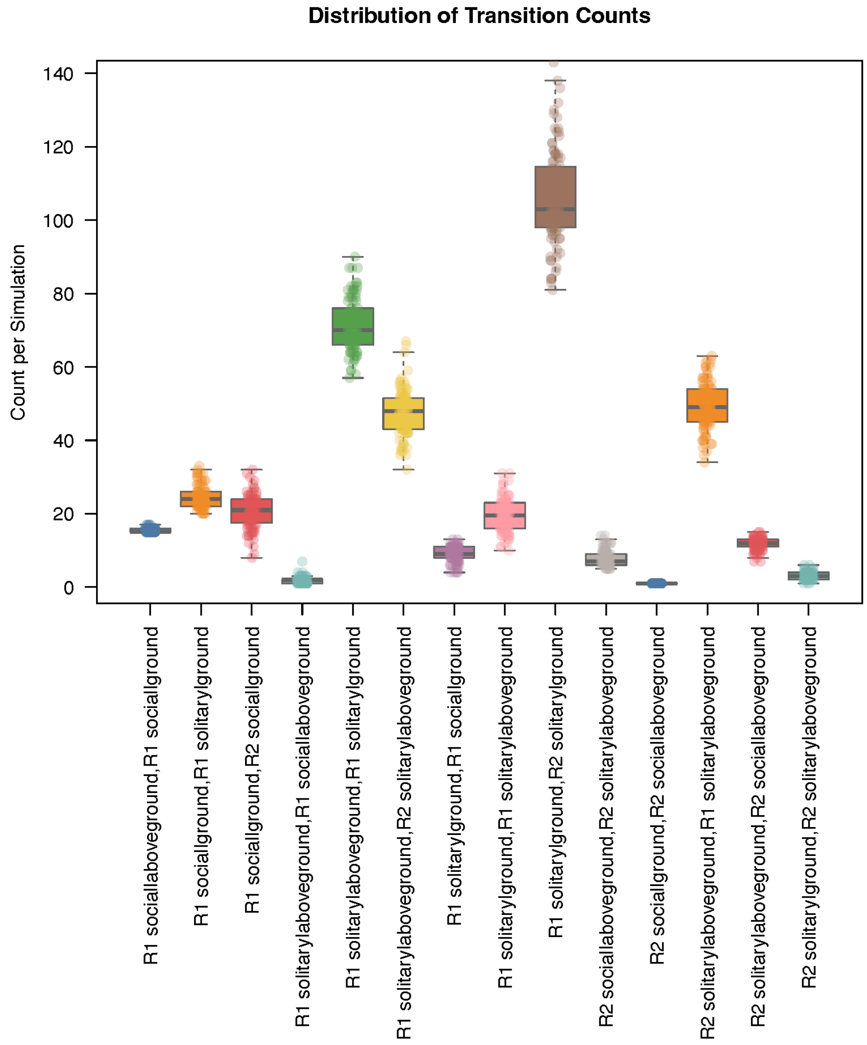

**Figure S4:** Summary of transition ages for 100 stochastic maps inferred from the corHMMdredge analysis. Note that some of the summarized transitions are between rate classes 1 (“fast”) and rate class 2 (“slow”).

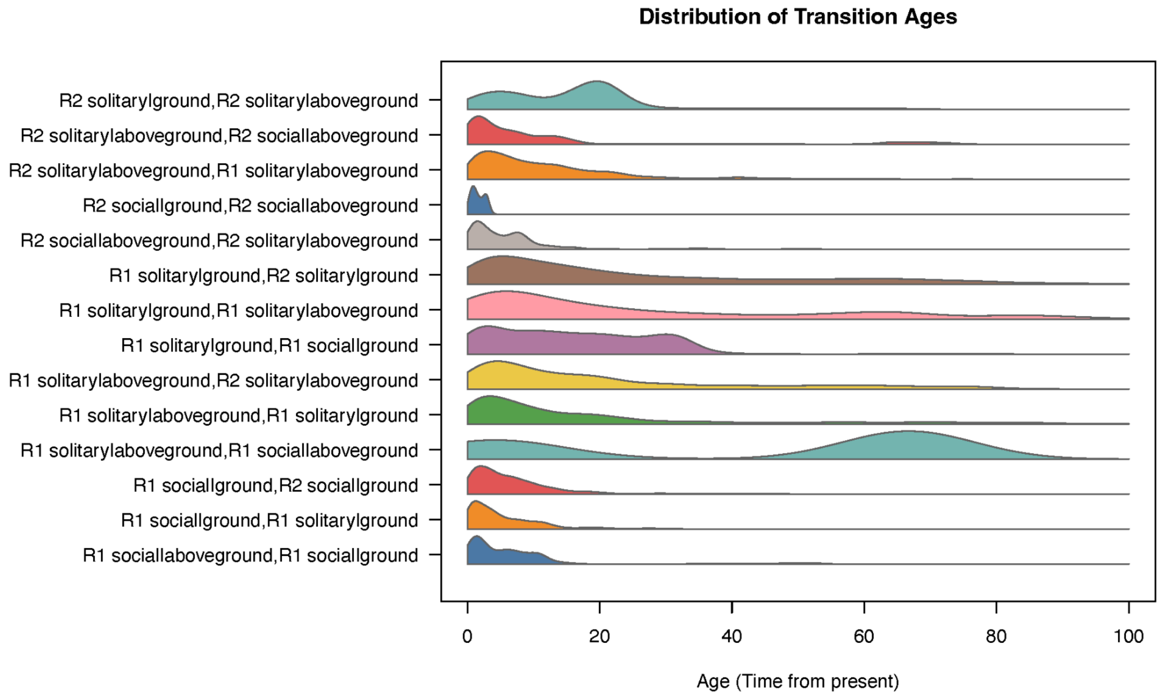

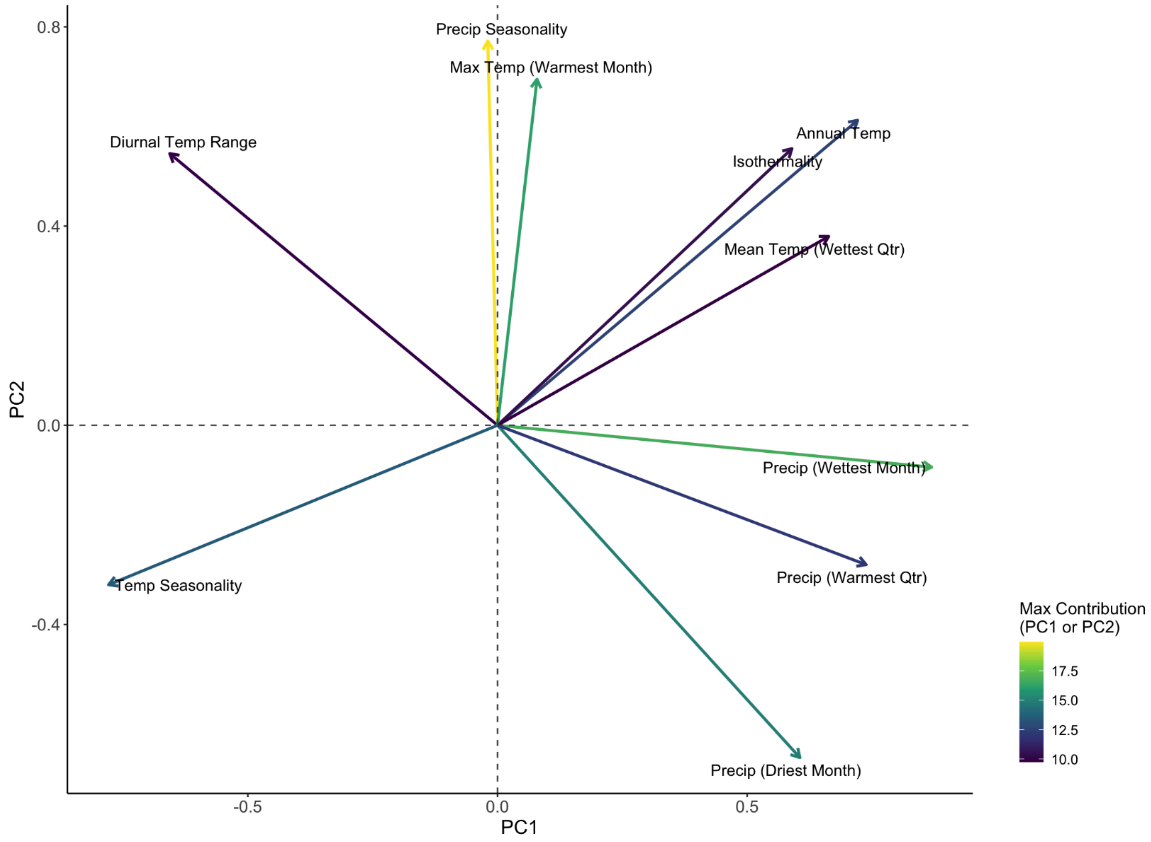


**Figure S5.** PCA loadings and contributions for top contributing variables (i.e., variables that contributed ≥ 9% to PC1 or PC2). Arrow direction represents the direction of the variable's influence on PC1 and PC2 (into which quadrant), arrow length represents the magnitude of the loading (i.e., how strongly the variable is associated with those axes), and arrow color represents the contribution (i.e., how much that variable contributes to explaining variance along PC1 or PC2).

### References

Henríquez-Piskulich, P., Hugall, A. F., & Stuart-Fox, D. (2024). A supermatrix phylogeny of the world's bees (Hymenoptera: Anthophila). *Molecular Phylogenetics and Evolution* 190: 107963. https://doi.org/10.1016/j.ympev.2023.107963

Karger, D. N., O. Conrad, J. Böhner, T. Kawohl, H. Kreft, R. W. Soria-Auza, N. E. Zimmermann, H. P. Linder, and M. Kessler. (2017). Climatologies at high resolution for the earth's land surface areas. *Scientific Data* 4(1): 170122. https://doi.org/10.1038/sdata.2017.122
