## Supplementary figures and images for "Sociality and nesting strategy shape the bimodal diversity gradient in bees"

### Figure S2

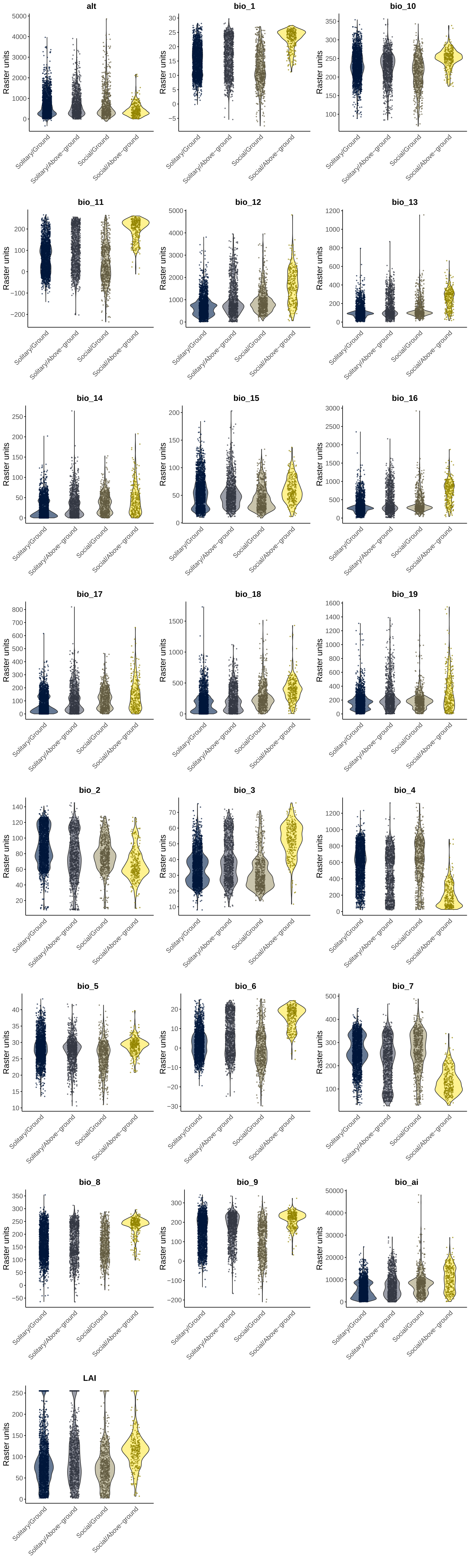
