## Supplementary material for "Sociality and nesting strategy shape the bimodal diversity gradient in bees": Table S3

Table S3: Dunn's post-hoc test results (Bonferroni-corrected) for climatic differences among bee trait groups. Arrows indicate whether the first group has significantly higher (↑) or lower (↓) values than the second group.

| **Climatic Variable** | **Comparison** | **Direction** | **Adj. p-value** | **Interpretation** |
|---|---|---|---|---|
| Mean annual temperature | Social aboveground – Social ground | ↑ social aboveground | $p < 0.0001$ | Among social bees, above-ground nesters occur in warmer climates than ground-nesters. |
| | Social aboveground – Solitary aboveground | ↑ social aboveground | $p < 0.0001$ | Among above-ground nesters, social bees occur in warmer areas than solitary bees. |
| | Social ground – Solitary aboveground | ↓ social ground | $p < 0.0001$ | Social ground-nesters occur in cooler areas than solitary above-ground. |
| | Social aboveground – Solitary ground | ↑ social aboveground | $p < 0.0001$ | Social above-ground bees occur in warmer areas than solitary ground-nesters. |
| | Social ground – Solitary ground | ↑ social ground | $p < 0.0001$ | Among ground-nesters, social bees occur in warmer areas than solitary bees. |
| | Solitary aboveground – Solitary ground | ↑ solitary aboveground | $p < 0.0001$ | Among solitary bees, above-ground nesters occur in warmer areas than ground-nesters. |
| Temperature seasonality | Social aboveground – Social ground | ↓ social aboveground | $p < 0.0001$ | Among social bees, above-ground nesters occur in less seasonal climates. |
| | Social aboveground – Solitary aboveground | ↓ social aboveground | $p < 0.0001$ | Among above-ground nesters, social bees are in less seasonal climates. |
| | Social ground – Solitary aboveground | ↑ social ground | $p < 0.0001$ | Social ground-nesting bees are in more seasonal climates than solitary above-ground. |
| | Social aboveground – Solitary ground | ↓ social aboveground | $p < 0.0001$ | Social above-ground nesting bees are in less seasonal climates than solitary ground-nesters. |
| | Social ground – Solitary ground | ↑ social ground | $p < 0.001$ | Among ground-nesters, social bees occur in more seasonal climates than solitary bees. |
| | Solitary aboveground – Solitary ground | ↓ solitary aboveground | $p < 0.0001$ | Among solitary bees, above-ground nesters occur in less seasonal environments than ground-nesters. |
| Annual precipitation | Social aboveground – Social ground | ↑ aboveground | $p < 0.0001$ | Among social bees, above-ground nesters occur in wetter environments than ground-nesters. |

*Continued on next page*

| Climatic Variable | Comparison | Direction | Adj. p-value | Interpretation |
|---|---|---|---|---|
| | Social aboveground – Solitary aboveground | ↑ social | $p < 0.0001$ | Among above-ground nesters, social bees occur in wetter areas than solitary bees. |
| | Social ground – Solitary aboveground | – (n.s.) | *n.s.* | No significant difference. |
| | Social aboveground – Solitary ground | ↑ aboveground social | $p < 0.0001$ | Social above-ground nesting bees occur in wetter areas than solitary ground-nesters. |
| | Social ground – Solitary ground | ↑ social | $p < 0.0001$ | Among ground-nesters, social bees occur in wetter environments than solitary bees. |
| | Solitary aboveground – Solitary ground | ↑ aboveground | $p < 0.0001$ | Among solitary bees, above-ground nesters occur in wetter environments than ground-nesters. |
| Precipitation seasonality | Social aboveground – Social ground | ↑ aboveground | $p < 0.0001$ | Among social bees, above-ground nesters occur in environments with greater precipitation seasonality. |
| | Social aboveground – Solitary aboveground | ↑ social | $p < 0.01$ | Among above-ground nesters, social bees occur in environments with greater precipitation seasonality. |
| | Social ground – Solitary aboveground | ↓ social ground | $p < 0.0001$ | Social ground bees occur in lower precipitation seasonality areas. |
| | Social aboveground – Solitary ground | ↑ aboveground social | $p < 0.01$ | Social above-ground bees occur in higher precipitation seasonality areas. |
| | Social ground – Solitary ground | ↓ social | $p < 0.0001$ | Among ground-nesters, social bees occur in lower precipitation seasonality areas. |
| | Solitary aboveground – Solitary ground | – (n.s.) | *n.s.* | No significant difference. |
