## Supplementary material for "Sociality and nesting strategy shape the bimodal diversity gradient in bees": Table S1

| **Species Name** | **Native Distribution** | **Regions of introduction** | **Reference** |
| --- | --- | --- | --- |
| *Anthidium manicatum* | Europe | NA, SA | Strange, JP, JB Koch, VH Gonzalez & L Nemelka. 2011. Global invasion by *Anthidium manicatum* (Linnaeus) (Hymenoptera: Megachilidae): assessing potential distribution in North America and beyond. *Biological Invasions* 13: 2115–2133. |
| *Anthophora villosula* | Asia | Maryland, US | `https://www.marylandbiodiversity.com/view/3139` |
| *Anthophora plumipes* (synonym of *A. villosula*) | — | Maryland, US | — |
| *Apis mellifera* | Europe, Asia, Africa | Americas, Australia, New Zealand | — |
| *Bombus terrestris* | Europe | SA, NA, Australia | Goulson, D. 2003. Effects of Introduced Bees on Native Ecosystems. *Annual Review of Ecology, Evolution, and Systematics* 34: 1–26. |
| *Bombus ruderatus* | Europe | NZ, SA | Goulson, D. 2003. Effects of Introduced Bees on Native Ecosystems. *Annual Review of Ecology, Evolution, and Systematics* 34: 1–27. |
| *Bombus hortorum* | Europe | NZ, Iceland | Goulson, D. 2003. Effects of Introduced Bees on Native Ecosystems. *Annual Review of Ecology, Evolution, and Systematics* 34: 1–28. |
| *Bombus lucorum* | Europe | NZ, Iceland | Goulson, D. 2003. Effects of Introduced Bees on Native Ecosystems. *Annual Review of Ecology, Evolution, and Systematics* 34: 1–29. |
| *Bombus subterraneus* | Europe | NZ | Goulson, D. 2003. Effects of Introduced Bees on Native Ecosystems. *Annual Review of Ecology, Evolution, and Systematics* 34: 1–30. |
| *Ceratina smaragdula* | Hawaii | India | Goulson, D. 2003. Effects of Introduced Bees on Native Ecosystems. *Annual Review of Ecology, Evolution, and Systematics* 34: 1–31. |

*Table Sx. (continued)*

| **Species Name** | **Native Distribution** | **Regions of introduction** | **Reference** |
|---|---|---|---|
| *Hylaeus punctatus* | Europe | NA, SA | Sheffield, C.S., Dumesh, S. & Cheryomina, M. 2011. *Hylaeus punctatus* (Hymenoptera: Colletidae), a bee species new to Canada, with notes on other non-native species. *Journal of the Entomological Society of Ontario* 142: 29–43. |
| *Megachile apicalis* | Europe | NA | Goulson, D. 2003. Effects of Introduced Bees on Native Ecosystems. *Annual Review of Ecology, Evolution, and Systematics* 34: 1–31. |
| *Megachile concinna* | Europe | California | Goulson, D. 2003. Effects of Introduced Bees on Native Ecosystems. *Annual Review of Ecology, Evolution, and Systematics* 34: 1–31. |
| *Megachile pusilla* | Palearctic | other parts of the world | Vossler, F.G. 2023. Pollen from urban flora in the ephemeral nests of solitary *Megachile* bees in three temperate and subtropical cities of Argentina. *Flora* 305: 152335. |
| *Megachile rotundata* | Eurasia | NA, Australia, NZ | Goulson, D. 2003. Effects of Introduced Bees on Native Ecosystems. *Annual Review of Ecology, Evolution, and Systematics* 34: 1–31. |
| *Megachile sculpturalis* | China, Japan | US, Europe | Goulson, D. 2003. Effects of Introduced Bees on Native Ecosystems. *Annual Review of Ecology, Evolution, and Systematics* 34: 1–32. |
| *Nomia melanderi* | North America | NZ | Goulson, D. 2003. Effects of Introduced Bees on Native Ecosystems. *Annual Review of Ecology, Evolution, and Systematics* 34: 1–33. |
| *Osmia caerulescens* | Europe | US | Goulson, D. 2003. Effects of Introduced Bees on Native Ecosystems. *Annual Review of Ecology, Evolution, and Systematics* 34: 1–34. |
| *Osmia cornifrons* | Japan | US | Goulson, D. 2003. Effects of Introduced Bees on Native Ecosystems. *Annual Review of Ecology, Evolution, and Systematics* 34: 1–35. |

| | | | |
|---|---|---|---|
| *Osmia ribifloris* | Southwestern US | Maine, US | Goulson, D. 2003. Effects of Introduced Bees on Native Ecosystems. *Annual Review of Ecology, Evolution, and Systematics* 34: 1–36. |
